# Plant pathogenic fungi hijack phosphate starvation signaling with conserved enzymatic effectors

**DOI:** 10.1101/2023.11.14.566975

**Authors:** Carl L. McCombe, Alex Wegner, Chenie S. Zamora, Florencia Casanova, Shouvik Aditya, Julian R. Greenwood, Louisa Wirtz, Samuel de Paula, Eleanor England, Sascha Shang, Daniel J. Ericsson, Ely Oliveira-Garcia, Simon J. Williams, Ulrich Schaffrath

## Abstract

Phosphate availability modulates plant immune function and regulates interactions with beneficial, phosphate-providing, microbes. Here, we describe the hijacking of plant phosphate sensing by a family of Nudix hydrolase effectors from pathogenic *Magnaporthe oryzae* and *Colletotrichum* fungi. Structural and enzymatic analyses of the Nudix effector family demonstrate that they selectively hydrolyze inositol pyrophosphates, a molecule used by plants to monitor phosphate status and regulate starvation responses. In *M. oryzae*, gene deletion and complementation experiments reveal that the enzymatic activity of a Nudix effector significantly contributes to pathogen virulence. Further, we show that this conserved effector family induces phosphate starvation signaling in plants. Our study elucidates a molecular mechanism, utilized by multiple phytopathogenic fungi, that manipulates the highly conserved plant phosphate sensing pathway to exacerbate disease.

**One-Sentence Summary:** A family of conserved enzyme effectors from pathogenic fungi manipulate plant phosphate sensing to promote infection.

## Main Text

Plant-microbe interactions range from beneficial to parasitic. Balancing the recruitment and support of symbiotic microbes, while maintaining the ability to defend against pathogens, is a major driver of plant evolution (*1*). The symbiosis between plants and arbuscular mycorrhizal fungi (AMF) predates the development of roots (*2*) and was likely instrumental in enabling land colonization by plants (*3*). Today, approximately 71% of vascular plants recruit AMF to access more phosphate and other mineral nutrients from the environment (*4*). In contrast to AMF, pathogenic fungi steal nutrients and constrain plant growth. Fungal diseases of important calorie crops threaten global food security by reducing crop yield (*5*). For example, *Magnaporthe oryzae* causes blast disease in major cereal crops including rice, wheat, and barley (*6*), resulting in annual food losses that could sustain hundreds of millions of people (*7*). Both symbiotic and pathogenic fungi secrete small proteins, called effectors, to optimize the host environment and support colonization.

Plant-AMF symbiosis is tightly regulated by the phosphate status of the plant (*8*). In eukaryotic cells, inositol pyrophosphates (PP-InsPs) signal phosphate availability through binding to SPX domains (*9*). When phosphate in plant cells is abundant, PP-InsP-bound SPX-domain proteins inhibit phosphate starvation response transcription factors (PHRs), thereby suppressing the expression of starvation induced genes (*10-13*). PHRs are conserved throughout land plants and green algae (*14*), and the regulation of plant-AMF symbiosis in both monocots and dicots is dependent on the PP-InsP/SPX/PHR signaling pathway (*15-18*). PHR activation also stimulates the expression of immune-suppressing genes and thereby inhibits the responsiveness of the plant immune system (*19,20*), indicating that plants prioritize symbiosis over defense during nutrient starvation. It is unknown whether any pathogenic fungi exploit the ancient and conserved phosphate-sensing pathway to promote plant infection. In this study, we demonstrate that a conserved family of Nudix (**Nu**cleoside-**di**phosphate linked to moiety-**X**) hydrolase effectors secreted by pathogenic fungi hydrolyze PP-InsPs. This function mimics phosphate depletion and hijacks the plant PHR signaling pathway. Gene deletion and complementation of *M. oryzae* Nudix effectors indicate that PP-InsP hydrolysis is essential for full virulence of the pathogen.

### A conserved Nudix hydrolase effector family promotes blast disease

Pathogenic *Magnaporthe* and *Colletotrichum* fungi possess effectors with putative Nudix hydrolase activity (Fig. 1A and table S1) (*21*). There are three predicted Nudix effector genes in *M. oryzae* (table S1) (*22*); two of these, named *MoNUDIX*, are identical in sequence and are highly upregulated during infection (*23*). To examine the role of Nudix effectors in cereal blast disease, we first utilized RNA interference (RNAi) to simultaneously lower the expression of both identical *MoNUDIX* genes (fig. S1, A and B). Silencing of *MoNUDIX* reduced blast disease symptoms in whole rice plant spray inoculation assays (fig. S1C). Furthermore, we report that silencing *MoNUDIX* enhances reactive oxygen species (ROS) accumulation, slows disease progression, and increases cell-wall autofluorescence (fig. S1, D to G). Collectively, the results indicate that *MoNUDIX* is important for *M. oryzae* virulence and host immune suppression. To corroborate the RNAi results and further characterize the Nudix effectors, we generated *MoNUDIX* deletion and complementation mutants in *M. oryzae* utilizing CRISPR/Cas9 genome editing (fig. S2). The deletion of both identical *MoNUDIX* gene copies (*M. oryzae*^ΔΔ*MoNUDIX*^) resulted in a drastic reduction in rice blast lesion size (Fig. 1, B and C). To assess whether the contribution of MoNUDIX to blast virulence was specific to rice infection, we also performed infection assays with barley and again observed significantly reduced lesion size with *M. oryzae*^ΔΔ*MoNUDIX*^ (Fig. 1, D and E). Our results were consistent across multiple rice and barley cultivars (Fig. 1, B and C, fig. S3, A and B), and microscopy analysis demonstrates a clear reduction in the growth of *M. oryzae*^ΔΔ*MoNUDIX*^ running hyphae during infection (fig. S3C). The two identical *MoNUDIX* genes likely function redundantly, as single deletion mutants exhibit only slight reductions in disease symptoms when compared to wild-type *M. oryzae* (fig. S4, A and B). *M. oryzae*^ΔΔ*MoNUDIX*^ displays normal vegetative growth, abiotic stress tolerance, conidia germination, appressorium formation, and infection following leaf wounding, which bypasses the requirement for an appressorium-mediated penetration (fig, S4, C to F). Collectively, our data indicate that MoNUDIX is important specifically for appressorium-mediated plant infection, this is consistent with the previously reported timing of Nudix effector gene induction during the late biotrophic growth stage of *M. oryzae, C. lentis*, and *C. higginsianum* (*21,23,24*).

**Fig. 1.**
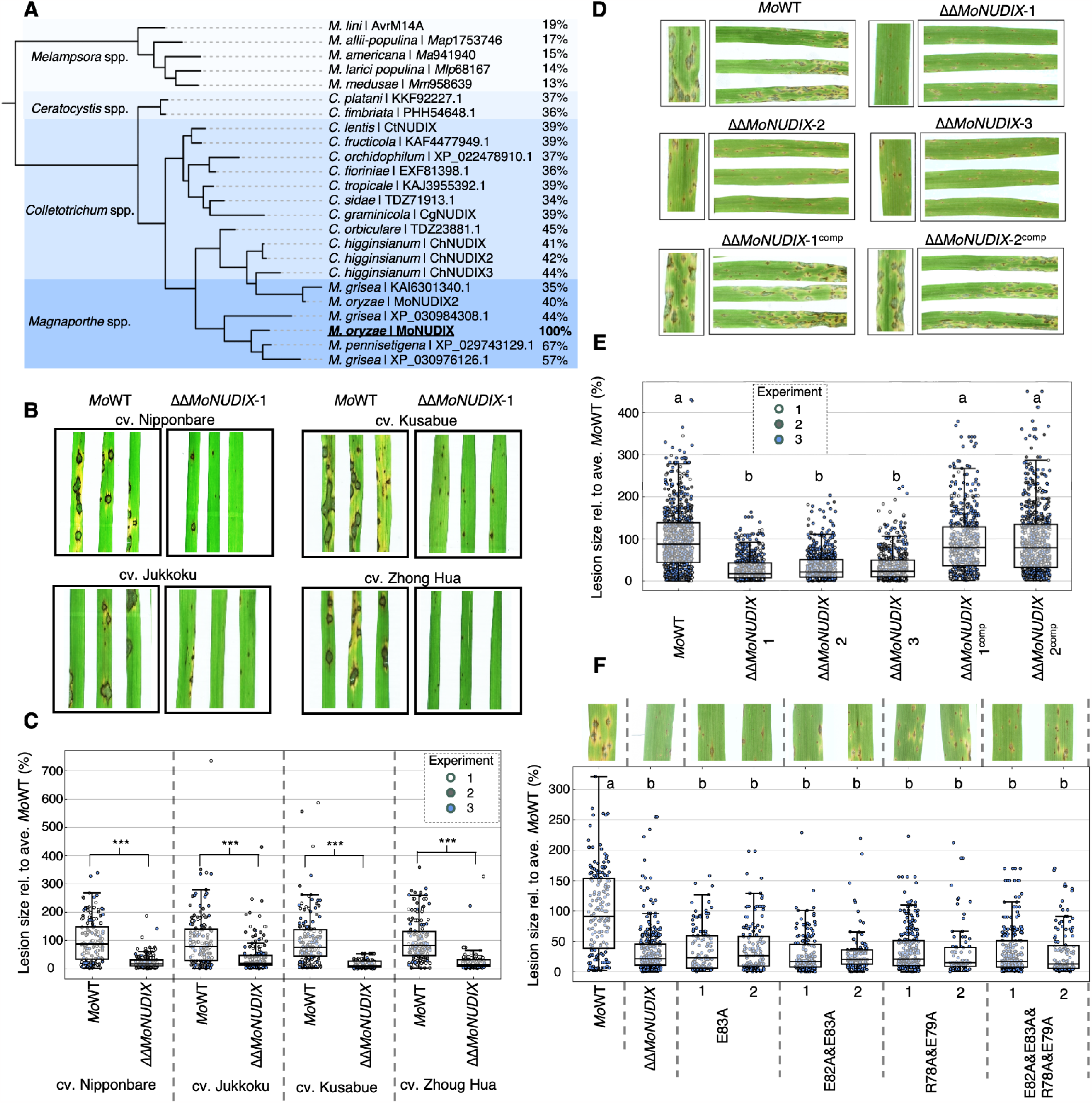
MoNUDIX is important for *Magnaporthe oryzae* virulence on rice and barley. **(A)** A phylogeny indicating the evolutionary relationships between Nudix hydrolase effectors from pathogenic fungi, the percentage value indicates amino acid sequence identity with MoNUDIX. **(B)** Rice cultivars Nipponbare, Jukkoku, Kusabue and Zhoug Hua were spray inoculated with *M. oryzae* isolate Guy11 (*Mo*WT) and *MoNUDIX* double gene deletion mutant ΔΔ*MoNUDIX*-1. Leaves were photographed six days post inoculation (dpi). **(C**) Boxplots summarize the results of three independent experiments, with the size of approximately 50 lesions measured across 5 rice leaves per treatment. Asterisks indicate treatments on the same cultivar with significantly different lesion sizes (Mann-Whitney U test, P < 0.001). (**D**) The barley cultivar Ingrid was inoculated with *Mo*WT, ΔΔ*MoNUDIX*-1, ΔΔ*MoNUDIX*-2, ΔΔ*MoNUDIX-3*, and MoNUDIX complementation mutants (ΔΔ*MoNUDIX*-1^comp^ and ΔΔ*MoNUDIX*-2^comp^). Leaves were photographed seven dpi. (**E**) Boxplots summarize the results of three independent experiments with approximately 150 lesions measured across 10 barley leaves per treatment. Letters depict significant differences between treatments (Kruskal-Wallis H test, Dunn’s post-hoc test, P < 0.01). (**F**) The barley cultivar Ingrid was inoculated with *Mo*WT, ΔΔ*MoNUDIX*-1, and ΔΔ*MoNUDIX*-1 complemented with MoNUDIX variants harboring substitution mutations in the Nudix box motif. Two independent complementation lines were tested for each variant. Example images at seven dpi are displayed. Boxplots summarize the results, the size of approximately 150 lesions were measured across 10 barley leaves. Letters depict significant differences between treatments (Kruskal-Wallis H test, Dunn’s post-hoc test, P < 0.01).

Nudix hydrolases possess a conserved sequence motif, GX_5_EX_7_REUXEEXGU, where U represents a hydrophobic amino acid and X is any amino acid (*25*). The first glutamate and arginine form a stabilizing salt bridge, while the remaining glutamates often bind divalent metal co-factors essential for enzymatic activity. To investigate the importance of MoNUDIX Nudix hydrolase activity, *M. oryzae*^ΔΔ*MoNUDIX*^ was complemented with either the wild-type effector, or effectors with various substitution mutations in the Nudix sequence motif. The expression of wild-type *MoNUDIX* successfully rescued the virulence of *M. oryzae*^ΔΔ*MoNUDIX*^, whereas expression of the mutant effector genes did not (Fig. 1, D to F). These data demonstrate that the Nudix motif and therefore hydrolytic activity is essential for the promotion of blast disease by MoNUDIX.

### *Magnaporthe* and *Colletotrichum* Nudix effectors are diphosphoinositol polyphosphate phosphohydrolases

Nudix hydrolases typically hydrolyze pyrophosphate bonds in molecules with a nucleoside diphosphate. To characterize the enzymatic activity and substrate specificity of the Nudix effectors, we determined the crystal structure of MoNUDIX. Structural similarity searches revealed that *Homo sapiens* diphosphoinositol polyphosphate phosphohydrolase 1 (*Hs*DIPP1) is remarkably similar to MoNUDIX in both overall structure and surface charge properties (Fig. 2A). *Hs*DIPP1, a well-characterized Nudix hydrolase, hydrolyzes diadenosine polyphosphates (Ap_n_As) (*26*), the protective 5′ mRNA cap (*27*), and PP-InsPs (*28*). Substrate screening with purified MoNUDIX protein demonstrated Nudix motif-dependent hydrolysis of 5-PP-InsP_5_ (Fig. 2B), while no activity was detected with Ap_n_As, mRNA caps, or other common substrates of Nudix hydrolases (fig. S5, A and B). Our results demonstrate that MoNUDIX is a selective diphosphoinositol polyphosphate phosphohydrolase *in vitro*. To understand the mode of PP-InsP binding to MoNUDIX, we modelled 5-PP-InsP_5_ into the MoNUDIX crystal structure via alignment with substrate bound *Hs*DIPP1 (*29*) (Fig. 2C). We identified basic amino acids likely required for PP-InsP binding (Fig. 2C); to confirm their involvement two lysines were mutated to glutamate (MoNUDIX^KKEE^). The MoNUDIX^KKEE^ protein demonstrated an approximately 70-fold reduction in inositol hexakisphosphate (InsP_6_) binding affinity as measured by micro-scale thermophoresis (fig. S5C) and was unable to hydrolyze 5-PP-InsP_5_ (Fig. 2D). The predicted PP-InsP binding site, including the basic amino acids, are conserved throughout the *Magnaporthe* and *Colletotrichum* Nudix effector family (Fig. 2, E and F), suggesting the conservation of substrate selectivity and enzymatic activity. To test this, we purified two homologs of MoNUDIX, one from *C. higginsianum* (ChNUDIX) and a second predicted *M. oryzae* Nudix effector (MoNUDIX2). Both ChNUDIX and MoNUDIX2 hydrolyze 5-PP-InsP_5_ and this activity is dependent on a Nudix motif glutamate (Fig. 2G). Conversely, AvrM14, a sequence-unrelated mRNA decapping Nudix hydrolase effector from the fungus *Melampsora lini* (*30*), does not hydrolyze 5-PP-InsP_5_ (fig S5D). All Nudix effectors used throughout this study were purified to homogeneity prior to *in vitro* characterization (fig. S6). Overall, structural analysis and enzymatic assays reveal that the *Magnaporthe* and *Colletotrichum* Nudix effectors are diphosphoinositol polyphosphate phosphohydrolases.

**Fig. 2.**
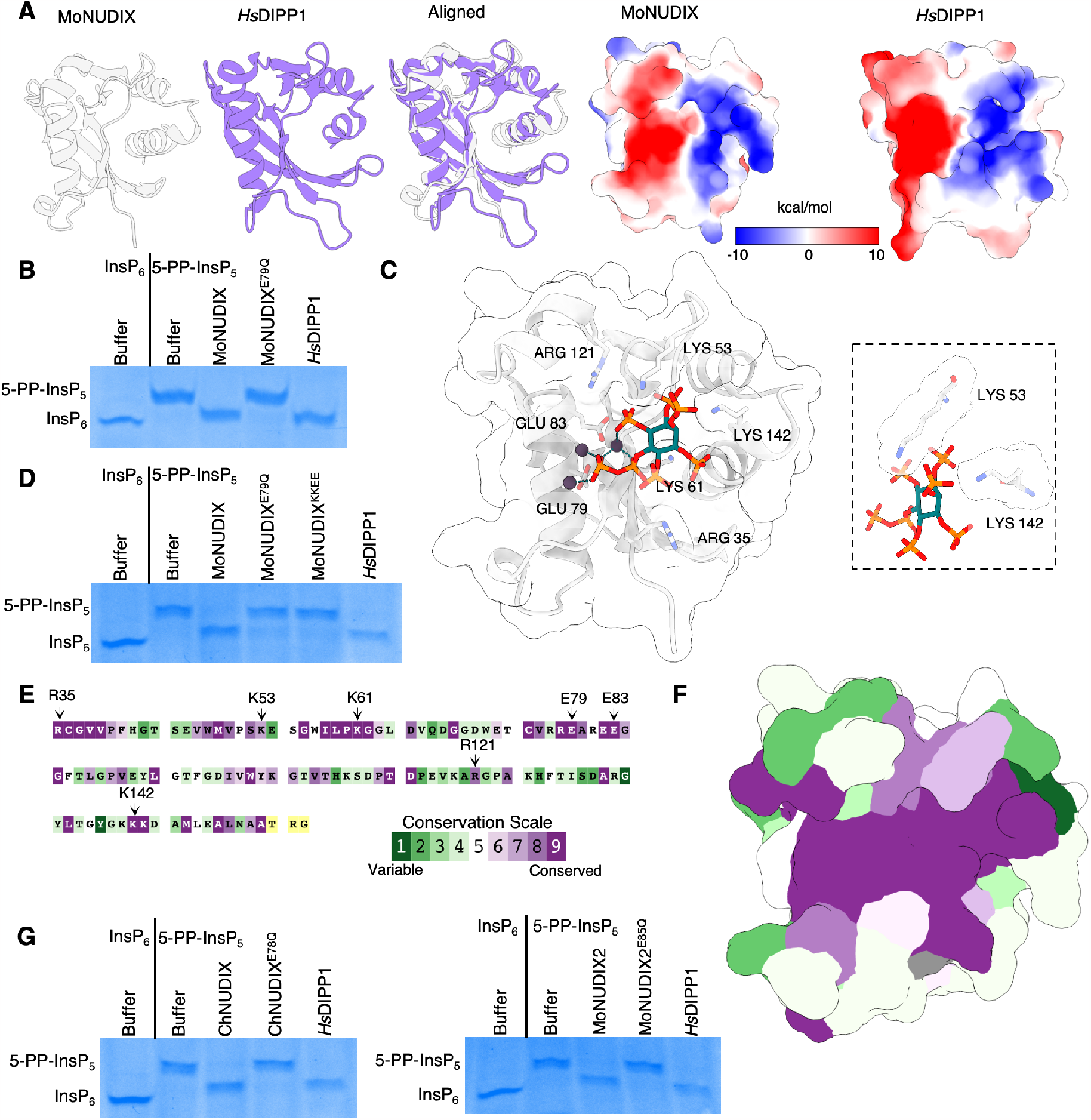
*Magnaporthe* and *Colletotrichum* Nudix effectors are diphosphoinositol polyphosphate phosphohydrolases. **(A)** MoNUDIX and *Homo sapiens* DIPP1 (PDB: 6WO7) (*26*) crystal structures superimposed to demonstrate their structural similarity despite low levels of sequence identity (25%). (Right) Both MoNUDIX and *Hs*DIPP1 demonstrate similar surface charge properties at the putative active site. (A) Purified MoNUDIX, MoNUDIX^E79Q^, and *Hs*DIPP1 were incubated with 5-PP-InsP_5_. Buffer alone was incubated with both InsP_6_ and 5-PP-InsP_5_. The reaction products were separated using a polyacrylamide gel and stained with toluidine blue. (**C**) MoNUDIX with Mg^2+^ and 5-PP-InsP_5_ docked into the crystal structure via alignment with *Hs*DIPP1 (PDB: 6WO7). The amino acids potentially important for Mg^2+^ and 5-PP-InsP_5_ binding are labelled. In the box are the two lysine amino acids selected for mutagenesis. (**D**) Inositol pyrophosphate hydrolysis assays demonstrate the importance of lysine 53 and 142. (**E**) The sequence of MoNUDIX used to determine the crystal structure, with colouring indicating amino acid conservation across homologous effectors. The amino acids labelled in (**C**) are indicated with arrows. (**F**) MoNUDIX with the protein surface coloured according to amino acid conservation across homologous effectors, demonstrating high conservation of the putative substrate binding site. (**G**) Inositol pyrophosphate hydrolysis assays with MoNUDIX homologs ChNUDIX and MoNUDIX2 demonstrates Nudix motif-dependent hydrolysis of 5-PP-InsP_5_.

### MoNUDIX localizes in the host cell cytoplasm during plant infection

*Magnaporthe oryzae* effectors can function within the host plant cell, or in the apoplastic space between the fungal cell wall and plant plasma membrane. The proteins involved in PP-InsP metabolism and signaling are intracellular. We therefore hypothesized that the Nudix effector family would function within the host plant cell during infection. We sought to identify the localization of MoNUDIX during plant infection using live-cell imaging techniques. First, we transformed *M. oryzae* with mRFP-tagged MoNUDIX controlled by the native promoter and determined that the effector co-localizes with the known cytoplasmic effector *Mo*Pwl2 in the biotrophic interfacial complex (BIC) (Fig. 3A). Using the native promoter was required for BIC localization, constitutive expression of MoNUDIX resulted in localization throughout the fungal hyphae (fig. S7). The BIC is the site of cytoplasmic effector translocation from the fungus into the host cell (*31, 32*), therefore our data suggest that MoNUDIX is a cytoplasmic effector. Treatment with brefeldin A (BFA), a potent inhibitor of Golgi trafficking (*33*), prevents the secretion of apoplastic but not cytoplasmic *M. oryzae* effectors (*32*). We demonstrate that the BIC localization of MoNUDIX is not influenced by BFA treatment, further indicating that MoNUDIX functions as a cytoplasmic effector (Fig. 3B). For concentration of cytoplasmic fluorescence and verification of MoNUDIX localization in the host cell cytoplasm, we utilized a gentle plasmolysis procedure. The MoNUDIX:mRFP signal is clearly observed in the host plant cell protoplasts following treatment with KNO_3_ (Fig. 3C). Overall, our results demonstrate that MoNUDIX localizes within the host cell cytoplasm during plant infection.

**Fig. 3.**
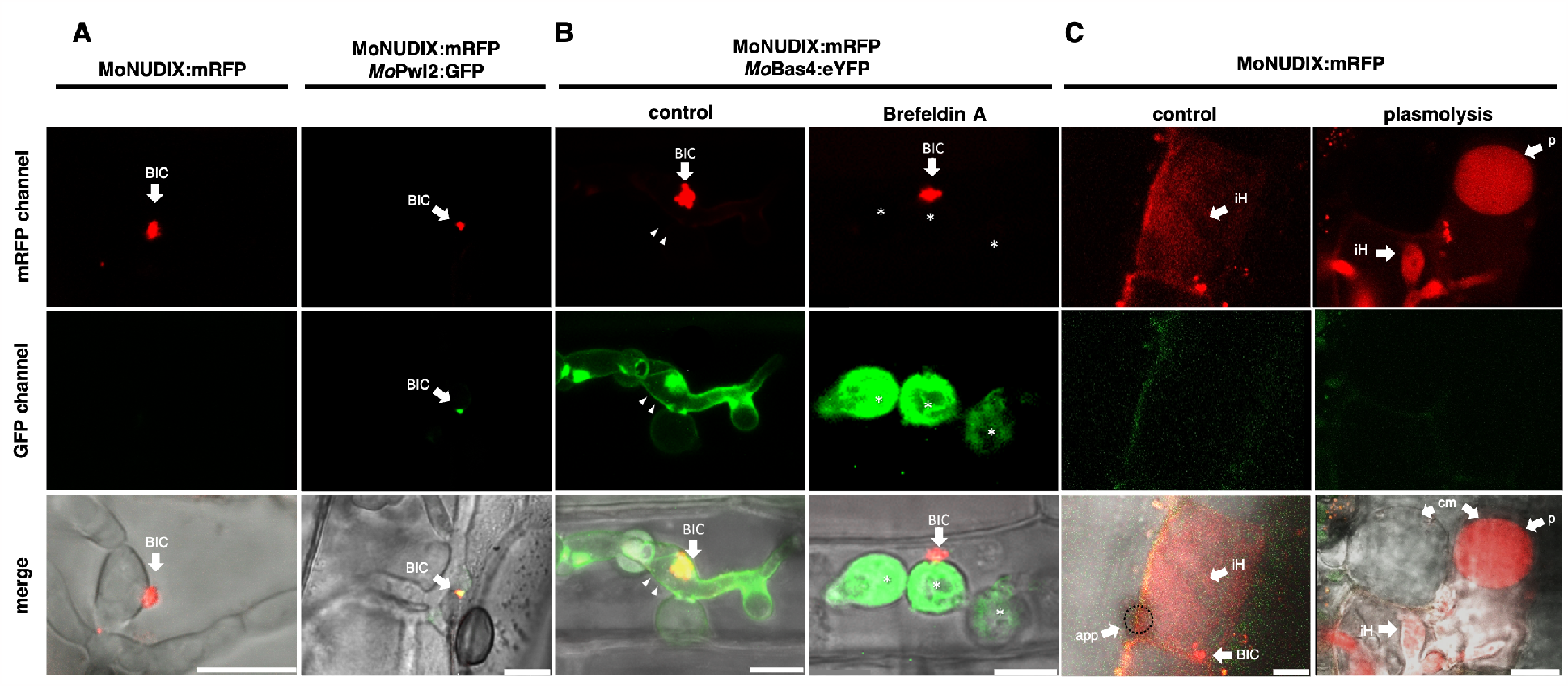
MoNUDIX localizes to the biotrophic interfacial complex and is secreted into the host cytoplasm during infection. Confocal laser-scanning microscopy images of barley and rice leaves infected with *Magnaporthe oryzae* expressing fluorescent protein-tagged effectors. (**A**) (Left) Barley leaves were inoculated with *M. oryzae* expressing MoNUDIX:mRFP. At 48 hours post inoculation (hpi), punctual accumulation of the mRFP fusion protein in the fungal hyphae was observed in the first infected cell, consistent with BIC localization. (Right) The cytoplasmic effector *Mo*Pwl2 *(27)* was co-expressed as a GFP-fusion protein demonstrating co-localization of MoNUDIX:mRFP and *Mo*Pwl2:GFP. (**B**) Rice leaves were inoculated with *M. oryzae* expressing MoNUDIX:mRFP and *Mo*Bas4:eYFP. *Mo*Bas4:eYFP (green) shows apoplastic localization outlining the invasive hypha (arrowheads). In the presence of BFA, *Mo*Bas4:eYFP (green) is retained in the fungal ER (asterisks), but MoNUDIX:mRFP remains BIC-localized (arrow), imaged with the same transformant at 3 h after exposure to BFA. **(C)** Secretion of MoNUDIX into the rice cell cytoplasm. Rice leaves were inoculated with *M. oryzae* expressing MoNUDIX:mRFP and analyzed 48 hpi. For concentration of the intracellular mRFP signal we used plasmolysis with 0.5 M KNO_3_ prior to imaging; BIC: biotrophic interfacial complex; iH: invasive hyphae; p: rice protoplast after plasmolysis; i: infected cell; cm: cell membrane of protoplast. Scale bar: 10 μm.

### The Nudix effector family activates plant phosphate starvation responses

A reduction in intracellular PP-InsP concentration, as occurs when phosphate is limited, leads to the activation of PHRs and phosphate starvation responses (PSRs). The depletion of PP-InsPs due to hydrolysis by the Nudix effectors should mimic localized phosphate starvation (fig. S8A). To determine if the Nudix effectors activate PHRs and PSRs in plants, we transiently expressed MoNUDIX, ChNUDIX, and mutated proteins without enzymatic activity (MoNUDIX^E79Q^ and ChNUDIX^E78Q^) in *Nicotiana benthamiana*. We selected two phosphate starvation induced (PSI) genes, *Nb*SPX1 and *Nb*PECP1, for gene expression analysis (*34, 35*). Both *Nb*SPX1 and *Nb*PECP1 have multiple PHR1-binding sequence (P1BS) elements in their promoter regions (fig. S8B). MoNUDIX and ChNUDIX significantly increased the abundance of *Nb*SPX1 and *Nb*PECP1 mRNA, when compared to MoNUDIX^E79Q^, ChNUDIX^E78Q^, and a no effector protein control (Fig. 4A). To investigate this further, we developed a rapid *in planta* screening method to monitor PHR activation. In our system, the RUBY reporter gene is controlled by a synthetic promoter with multiple P1BS elements, we named the promoter/reporter construct PSI:RUBY (fig. S8C). This system allowed us to screen six Nudix effector family members along with their corresponding Nudix motif mutants and other negative controls (nanoluciferase and AvrM14). We report that all six Nudix effectors significantly increase the expression of the RUBY reporter when compared to the corresponding Nudix motif mutant proteins and the negative controls (Fig. 4, B and C). Additionally, MoNUDIX^KKEE^ expression results in reduced RUBY expression when compared to wild-type MoNUDIX, consistent with the reduced InsP_6_ binding and a lack of 5-PP-InsP_5_ hydrolysis observed *in vitro* (Fig. 4, B and C). We also observe similar results when swapping the synthetic P1BS promoter with the 1 kb promoter region from *Nb*PECP1 (fig. S8, D and E). For all proteins, accumulation in *N. benthamiana* leaf tissue was detected via immunoblotting (fig. S8F). Collectively, our data demonstrate that the enzymatic activity of the Nudix effectors activates PHRs and PSI gene expression in *N. benthamiana*.

**Fig. 4.**
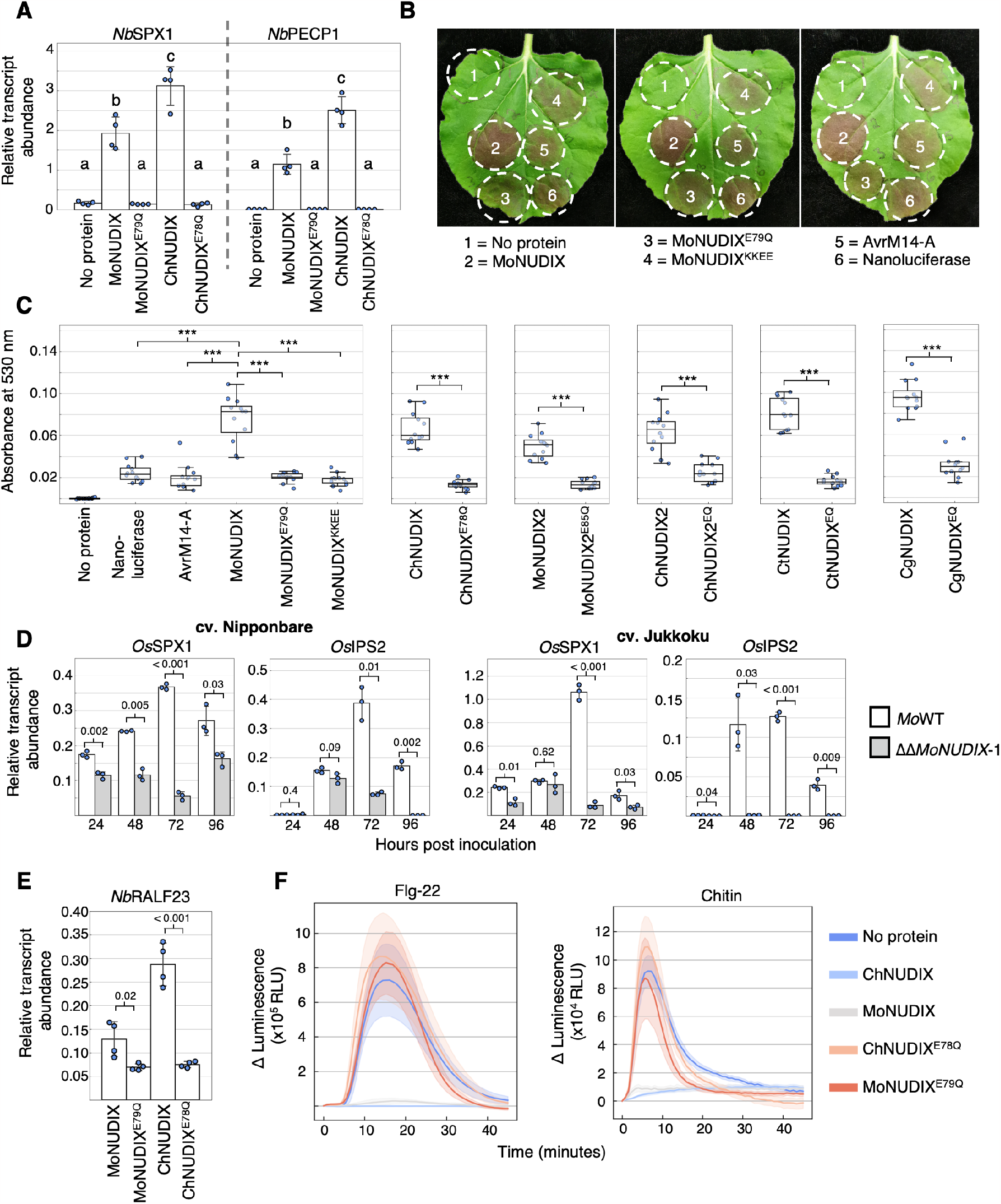
The Nudix effector family activates plant phosphate starvation responses. (**A**) The transcript abundance of *Nb*SPX1 and *Nb*PECP1 in *N. benthamiana* expressing Nudix effectors, or leaves transformed with an empty vector control (no protein). Values are mean expression ± SD (n = 4) relative to the reference genes, letters indicate significantly different groups (one-way ANOVA, Tukey’s HSD, P < 0.001). (**B**) Visible RUBY production in leaves co-transformed with PSI:RUBY and Nudix effectors or controls. (**C**) The absorbance of extracts from leaves co-transformed with PSI:RUBY and Nudix effectors or controls (n = 12), asterisks indicate significant differences between treatments (one-way ANOVA, Tukey’s HSD, *** P <0.001). (**D**) Rice cultivars Nipponbare and Jukkoku were inoculated with *M. oryzae* isolate Guy11 (*Mo*WT) and *M. oryzae*^ΔΔ*MoNUDIX*-1^. *Os*SPX1 and *Os*IPS2 transcript abundance was calculated relative to *Os*ACTIN at four timepoints throughout the infection process, values are mean ± SD (n = 3), significant differences between *Mo*WT and ΔΔ*MoNUDIX*-1 treatments were identified by an independent samples t test, p-values are listed. (**E**) The expression of *Nb*RALF23 in *N. benthamiana* leaves expressing Nudix proteins (as labelled). Values are mean ± SD (n=4) relative to the reference genes, with p-values from an independent samples t-test displayed. (**F**) Reactive oxygen species production in *N. benthamiana* expressing Nudix effectors, or leaves transformed with an empty vector control (no protein), following exposure to flg-22 (left) or chitin (right). Values are mean (solid line) ± SEM (shaded area) (n = 8).

To determine if MoNUDIX induces PSRs during rice infection, we compared the mRNA levels of two PSI rice genes (*Os*SPX1 and *Os*IPS2) in two different rice cultivars throughout the infection process with either wild-type *M. oryzae* or *M. oryzae* ^ΔΔ*MoNUDIX*^. At 72 hours post inoculation the expression of both *Os*SPX1 and *Os*IPS2 is significantly upregulated in rice infected with wild-type MoNUDIX when compared to *M. oryzae* ^ΔΔ*MoNUDIX*^, suggesting that MoNUDIX promotes the expression of PSI genes in rice (Fig. 4D).

In *A. thaliana*, phosphate starvation induces the expression of immunosuppressant rapid alkalization factor (RALF) genes with P1BS elements in their promoter region (for example *At*RALF23) (*20, 36*). This reduces reactive oxygen species (ROS) production triggered by immune elicitors (*20, 36*). Similarly, in *N. benthamiana* the enzymatic activity of the Nudix effectors induces the expression of an *At*RALF23 homolog (designated *Nb*RALF23) and prevents the immune-activated ROS burst (Fig. 4, E and F). Overall, our results demonstrate that the diphosphoinositol polyphosphate phosphohydrolase effectors can activate phosphate starvation responses, which likely results in the suppression of plant immunity.

## Discussion and conclusion

Phosphate is an essential but often limiting nutrient for plant growth, with phosphate-starved plants actively recruiting soil microbes to improve phosphate acquisition. Plant phosphate status is important for the regulation of multiple plant-microbe interactions (*37-42*). Here, we describe the molecular basis for phosphate status manipulation by pathogenic fungi. Our findings provide direct evidence for the hijacking of symbiosis-facilitating mechanisms by pathogenic fungi to promote disease.

The deletion of *MoNUDIX* genes in *M. oryzae* resulted in impaired plant colonization and fungal growth. The infection phenotype of *M. oryzae*^ΔΔ*MoNUDIX*^ is analogous to phr2-mutant rice inoculated with AMF (*17*) and suggests that PHR-activation can enhance both beneficial and pathogenic infection in rice. Like AMF, *Colletotrichum tofieldiae* promotes host plant growth by providing phosphate under starvation conditions (*38*). We were unable to identify Nudix effectors in *C. tofieldiae*, and their absence may be required to ensure *C. tofieldiae* growth remains appropriately regulated by plant phosphate status. Conversely, multiple pathogenic *Colletotrichum* species have Nudix effectors that we demonstrate activate starvation responses. Three *C. higginsianum* Nudix effectors (two copies of *ChNUDIX*, one copy of *ChNUDIX2*) are clustered together on a mini-chromosome (chromosome 11) and are highly upregulated early in plant infection (*24*). Similar to *M. oryzae*^ΔΔ*MoNUDIX*^, C. *higginsianum* mutants lacking chromosome 11 display normal vegetative growth and can successfully penetrate host plant cells but demonstrate inhibited disease progression (*43*). There are only 8 predicted effector genes on chromosome 11 (*44*), and therefore it is likely that Nudix effectors are also important for the pathogenicity of *C. higginsianum*.

In addition to their central role in phosphate homeostasis, PP-InsPs are co-factors for the receptors sensing the phytohormones auxin and jasmonate (*45-48*). Phosphate status and jasmonate signaling are intricately linked in plants. For example, jasmonate treatment stimulates PP-InsP synthesis (*49*), and PHR-activation enhances jasmonate production and signaling (*50, 51*). By manipulating intracellular PP-InsP levels, the Nudix effectors may influence jasmonate and auxin signaling in their host plants in addition to inducing starvation responses.

Based on our data, we propose the following model describing the function of the *Magnaporthe* and *Colletotrichum* Nudix hydrolase effector family (Fig. 5). First, the effectors are translocated into their respective host plant cells. Once inside, they function as PP-InsP hydrolase enzymes, effectively uncoupling PHR activation from intracellular phosphate availability. This induces a plethora of transcriptional changes to promote phosphate acquisition and suppress immune responses, ultimately promoting disease.

**Fig. 5.**
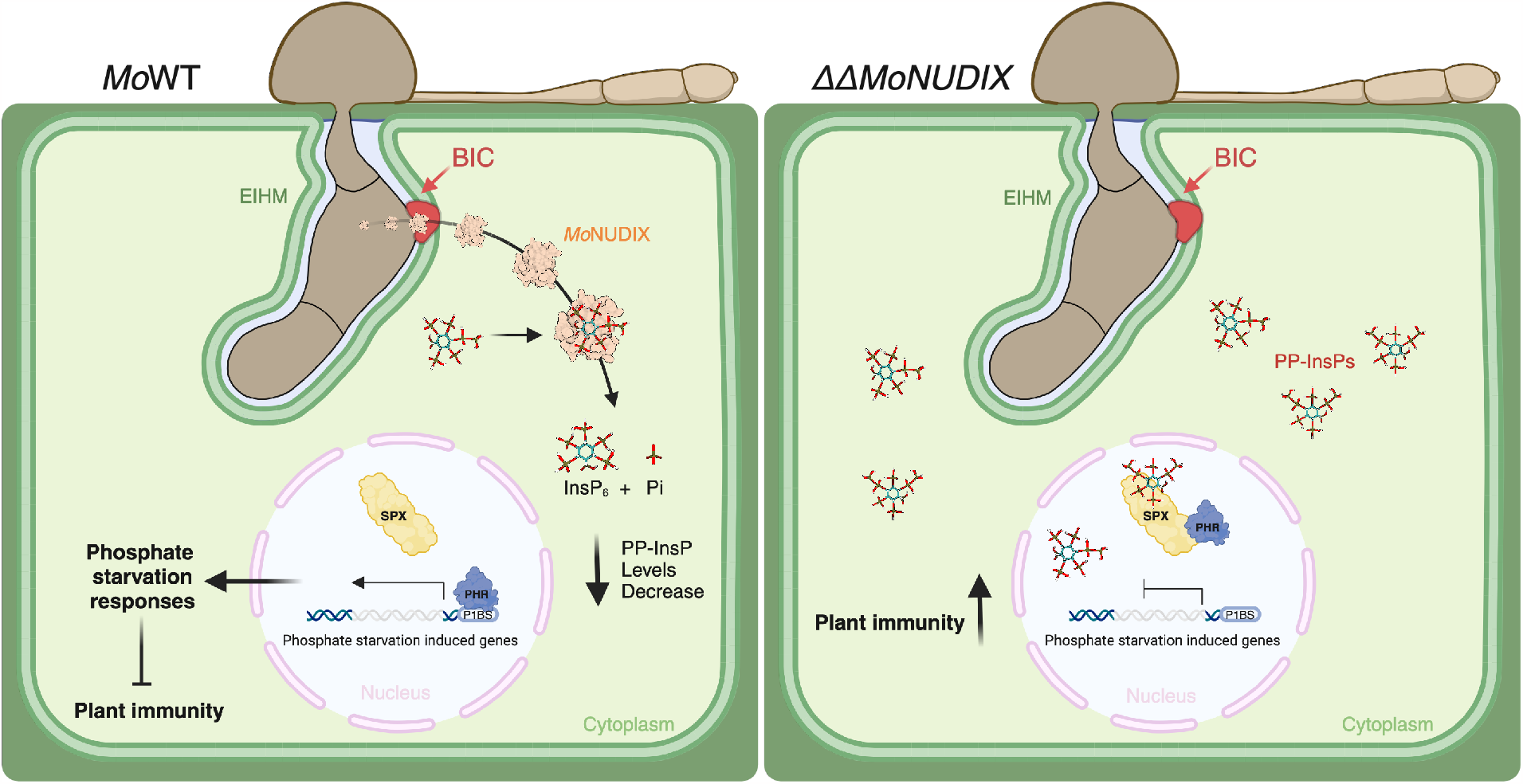
A model for the virulence function of the *Magnaporthe* and *Colletotrichum* Nudix hydrolase effectors. (Left) In wild-type *M. oryzae* (*Mo*WT) MoNUDIX is secreted from the invading fungus into the host cell cytoplasm. MoNUDIX functions as an enzyme, hydrolyzing inositol pyrophosphate (PP-InsP) signaling molecules into inositol hexakisphosphate (InsP_6_) and phosphate (Pi). The decrease in PP-InsP concentration prevents SPX-mediated inhibition of PHRs, resulting in the transcription of phosphate starvation inducible genes. Phosphate starvation responses are therefore activated, including those which suppress plant immune responses. (Right) Without MoNUDIX, the ability of *M. oryzae*^ΔΔ*MoNUDIX*^ to stimulate PP-InsP hydrolysis is compromised. Available PP-InsPs are detected by SPX domains, resulting in the binding of SPX domains to PHRs. This prevents PHRs from binding to the P1BS elements in phosphate starvation induced genes. In the absence of phosphate starvation signaling, plant immune responses are prioritized.

## Acknowledgments

We thank the Plant Services Team at the Australian National University for providing *N. benthamiana* seedlings, and all the Williams lab members for helpful discussion. The authors acknowledge the use of the ANU crystallization facility. The authors also acknowledge the use of the Australian Synchrotron MX facility and thank the staff for their support. This research was undertaken in part using the MX2 beamline at the Australian Synchrotron, part of ANSTO, and made use of the Australian Cancer Research Foundation (ACRF) detector. Fig. 5 and S8A were created with BioRender.com.

## Funding

Australian Institute of Nuclear Science and Engineering Ltd. Postgraduate Research Award (CLM)

Australian Government Research Training Programme Stipend (CLM)

ANU Future Scheme 35665 (SJW)

ARC Future Fellowship FT200100135 (SJW)

Louisiana Board of Regents grant #LEQSF (2022-24)-RD-A-01 (EOG)

RWTH Aachen University scholarships for Doctoral Students (AW, FC, LW)

Funded by the Deutsche Forschungsgemeinschaft (DFG, German Research Foundation) – 410278620 (US, AW)

## Author contributions

Conceptualization: CLM, AW, SJW, US

Methodology: CLM, AW, SA, JRG, EOG, SJW, US

Investigation: CLM, AW, CSZ, FC, SA, JRG, LW, SdP, EE, SS, EOG

Formal Analysis: CLM, AW, CSZ, FC, SA, DJE, EOG, SJW

Visualization: CLM, AW, CSZ, FC, SA, EOG

Funding acquisition: EOG, SJW, US

Project administration: CLM, AW, SJW, US

Supervision: CLM, AW, EOG, SJW, US

Writing – original draft: CLM, AW

Writing – review & editing: CLM, AW, FC, SA, JRG, LW, SS, EE, DJE, EOG, SJW, US

## Data and materials availability

Where possible all materials generated in this study will be made available upon request to the corresponding authors. The data that support the MoNUDIX protein structure described in this study are openly available under accession 8SXS at the PDB. All other data are available in the main text or supplementary materials.

## Supplementary Materials

Materials and Methods

Figs. S1 to S8

Tables S1 to S6

References (*52-88*)

