## supplementary materials for "Plant pathogenic fungi hijack phosphate starvation signaling with conserved enzymatic effectors"

**The PDF file includes:**

Materials and Methods
Figs. S1 to S8
References 52 - 88

**Other Supplementary Materials for this manuscript include the following:**

Tables S1 to S6

### Materials and Methods

Plant growth and *Magnaporthe oryzae* infection conditions

*Nicotiana benthamiana* plants were grown in a controlled environment at 25 °C under a 16-hour/8-hour light/dark photoperiod. Fully expanded leaves from 5-week-old plants were used in Agrobacterium-mediated transient gene expression experiments to generate qPCR, RUBY promoter/reporter and ROS burst assay results.

Rice (*O. sativa*) plants for RNAi and BFA-treatment protein localization experiments were grown in Baccto Top Soil (Michigan Peat Co., Houston, Texas) in a Caron 7301-50 Plant Growth Chamber with equal numbers of fluorescent lamps (Philips ED37, 400 W). At rice seedling height, ~1 m from the bulbs, light ranged in intensity from 600 to 1,200 µmol $m^{-2}$ $s^{-1}$. Plants were grown at ~70 % relative humidity under a daily cycle of 12 h of light at 28 °C and 12 h of darkness at 24 °C. At 2 weeks, plants were fertilized with Jack's Professional Peat Lite 20-10-20 Fertilizer (#77860; JR Peters, Inc.; Allentown, PA). Whole plant infection assays for RNAi experiments were performed by spray inoculation of 2–3-week-old rice plants as previously described (*52*).

Rice and barley used for the plasmolysis localization experiments and for the characterization of infection phenotypes with *M. oryzae MoNUDIX* gene deletion and complementation mutants. Barley plants were cultivated in plant-growth chambers at a temperature of 18 °C, relative humidity of 60 %, and a 16/8 h light-dark cycle with a light intensity of 210 µmol/$m^2$s in ED73-soil (Balster Einheitserdewerk GmbH, Germany), as previously described (*53*). After 7 days the primary leaves of the plants were inoculated with *M. oryzae* (10000 conidia $mL^{-1}$) using a compressed air atomizer. Subsequently, the plants were kept in the dark at 24 °C and 100 % relative humidity for 24 hours before being transferred to growth chamber conditions. Rice plants were cultivated in P-soil (Balster Einheitserdewerk GmbH, Germany) at 24 °C in a growth chamber with a 15/9 h light-dark cycle, 80 % relative humidity, and a light intensity of 300 µmol/$m^2$s. Rice plants were spray inoculated after 12 days with 20000 conidia $mL^{-1}$. Subsequently, the plants were kept in the dark at 24 °C and 100 % relative humidity for 24 hours before being transferred to growth chamber conditions. Infected rice leaves were evaluated at 5 dpi, barley leaves at 6 dpi. Images were captured using a Nikon D3500 camera. Disease symptoms were quantified by determining the size in pixels of each symptom using ImageJ software. All reported values in the manuscript are relative to the average lesion size for the wild-type *M. oryzae* isolate Guy11.

Fungal cultivation

For RNAi and BFA-treatment protein localization experiments, fungal strains (*M. oryzae*) were stored on dried filter papers at −20˚C and cultured on rice bran agar plates at 25°C for up to 2 weeks under continuous light in Percival Scientific (Model CU-36L5) tissue culture incubators equipped with one half fluorescent lights (FT20T12/cw, 20W) and one-half black lights (FT20T12/BL, 20W).

For all other experiments, the original *M. oryzae* strain Guy11 (*Mo*WT) was provided by Dr. D. Tharreau from CIRAD in Montpellier, France. Fungi were cultivated on oatmeal agar, which consisted of 20 g/L agar, 2 g/L yeast extract, 10 g/L starch, and 30 g/L oat flakes. Alternatively, they were grown on potato dextrose agar (PDA) at a temperature of 23 °C in the absence of light. Conidia were obtained as described in (*53*) by cultivating the fungi placed under fluorescent tubes emitting blacklight (310 to 360 nm) with a 16/8 h light/dark cycle and at a constant temperature of 26 °C for 7 days. The concentration of conidia was adjusted in a solution containing 1 g/L gelatin and 0.5 ml/L Tween 20.

Plasmid construction for gene expression in *E. coli* and *N. benthamiana*

All effector gene sequences had their signal peptide predicted using SignalP 6.0 (*54*) and removed, were codon optimised for *E. coli*, and ordered from Integrated DNA technologies, Inc IDT® as double-stranded DNA fragments. The DNA fragments were cloned into either a modified pOPIN plasmid (*55*) with a T7 promoter and a 6xHIS 3C protease site N-terminal tag if used for protein expression in *E. coli*, or a level 1 MoClo plasmid (*56*) along with a 35 S promoter, Omega 5′ UTR translational enhancer, 3xHA N-terminal tag, and octopine synthase terminator if used for expression in *N. benthamiana*. The sequences of all proteins expressed in *E. coli* and *N. benthamiana* in this study are listed in **Table S2** and **Table S3**. All level 1 MoClo genes were subsequently inserted into the level 2 acceptor plasmid pICSL4723 before transformation into *Agrobacterium tumefaciens* (GV3101). The RUBY polyprotein gene sequence, and all gene promoter sequences used in this study (listed in **Table S4**) were ordered from Integrated DNA technologies, Inc IDT® as double-stranded DNA fragments and cloned into level 0 MoClo plasmids (*56*), before being used to create level 1 and level 2 plasmids. *Entamoeba histolytica* IP6KA and *Homo sapiens* DIPP1 were ordered as *E. coli*-codon optimised double-stranded DNA fragments and cloned into the modified pOPIN (*53*) plasmid with a T7 promoter and a 6xHIS 3C protease site N-terminal tag. All gene sequences had any BbsI and BsaI cleavage sites removed without altering protein sequence to enable GoldenGate assembly.

Protein expression and purification

AvrM14-A, *Hs*Nudt16, *At*Nudx7, and *At*Nudx7$^{E154Q}$ proteins were purified previously (*30*). *Eh*IP6KA and *Hs*DIPP1 were expressed in *E. coli* BL21 (DE3) cells using ZYM-5052 autoinduction media (*57*). Cells were grown by continuous shaking at 37 °C until the $OD_{600nm}$ reached 0.6 – 0.8. The temperature was then dropped to 18 °C and cells were incubated with shaking for another 18 hours before harvesting via centrifugation. MoNUDIX, MoNUDIX2, and ChNUDIX proteins (wild-type and mutants) were expressed in *E. coli* Shuffle® cells grown in Terrific Broth. Cells were grown by continuous shaking at 30 °C until the $OD_{600nm}$ reached 0.6 – 0.8. The temperature was then dropped to 16 °C, IPTG was added to a final concentration of 200 µM, and incubation with shaking continued for another 18 hours before harvesting via centrifugation. Following centrifugation, all cell pellets were resuspended in lysis buffer (50 mM HEPES pH 8.0 (MoNUDIX, MoNUDIX$^{E79Q}$, MoNUDIX$^{KKEE}$), pH 7.5 (MoNUDIX2, MoNUDIX2$^{EQ}$ ChNUDIX, ChNUDIX$^{E78Q}$), or pH 7.0 (*Hs*DIPP1, *Eh*IP6KA), 150 mM NaCl, 1 mM PMSF, 1 µg $ml^{-1}$ DNase, and 1 mM DTT (DTT only included for *Eh*IP6KA and *Hs*DIPP1)). All cells were lysed using sonication and cellular debris pelleted by centrifugation. The resulting supernatant was applied to a 5 mL HisTrap FF crude column (Cytiva, Marlborough, Massachusetts). To remove loosely bound proteins, the column was washed with the lysis buffer without PMSF or DNase containing 30 mM imidazole. The remaining bound proteins were eluted with a continuous gradient of imidazole from 30 mM to 250 mM over 10 minutes, using an Äkta pure chromatography system. Fractions were analysed by Coomassie-stained SDS-PAGE and fractions containing the protein of interest were pooled, dialysed to remove the imidazole, and incubated with recombinant 6xHis-tagged 3C protease overnight at 4 °C (except *Eh*IP6KA, which was stored following dialysis without 3C protease incubation). The protein of interest was separated from any uncleaved protein, the fusion tag, and 3C protease by immobilized metal affinity chromatography and purified further by size-exclusion chromatography (SEC) using either a HiLoad 16/600 Superdex® 75 pg or a HiLoad 26/600 Superdex® 75 pg column pre-equilibrated in buffer (10 mM HEPES (pH same as in corresponding lysis buffer), and 150 mM NaCl). After

SEC, fractions containing the protein of interest were identified using SDS-PAGE and concentrated using Amicon® Ultra Centrifugal filters (Merck, Darmstadt, Germany) before storage at -80 °C.

Putative Nudix hydrolase effector identification and phylogenetic tree construction

To identify putative Nudix hydrolase effectors homologous to MoNUDIX, the NCBI protein database (*58*) was searched using blastp with the protein sequence of MoNUDIX (default parameters; word size = 5, expect threshold = 0.05). Any identified hits without a predicted signal peptide or Nudix hydrolase domain were filtered out by screening the sequences using SignalP6.0 (*54*) and InterProScan (*59*). To reduce the length of the list while retaining sequence diversity, if two or more sequences shared > 95% sequence identity, only one sequence was selected at random to remain in the list presented in **Table S1**. PhyML (*60*) (version 3.3) was used to estimate a maximum-likelihood phylogeny with selected protein sequences from this list, along with previously identified fungal Nudix effectors (*30*). The resulting phylogeny was visualized using iTOL (*61*) (version 6.7.6).

Transient gene expression via agrobacterium infiltration

*Agrobacterium tumefaciens* (GV3101) with the desired plasmid was suspended in infiltration buffer (10 mM MES pH 5.6, 10 mM MgCl2 and 200 µM acetosyringone) to an optical density at 600 nm ($OD_{600nm}$) of 0.5 (for RUBY co-infiltrations) or 1.0 (for qPCR and the ROS burst assay). For co-infiltrations, *A. tumefaciens* (GV3101) with a RUBY promoter/reporter plasmid was added to the infiltration buffer to an $OD_{600nm}$ of 0.5. The final combined $OD_{600nm}$ was 1.0 for all infiltrations. All cultures were incubated in the dark at 28 °C with 220 rpm shaking for 2 to 3 hours before syringe-infiltration into *N. benthamiana* leaves. Infiltrated plants were kept in the same growing conditions as before infiltration.

ROS burst assay

Measurement of ROS was completed as described previously with some minor modifications (*62*). In brief, *N. benthamiana* leaf discs (4 mm diameter) were floated on water overnight in a 96-

well plate. The water was replaced with an elicitor solution (200 µM luminol, 20 µg ml−1 horseradish peroxidase and 100 nM flg-22 or 5 µg $mL^{-1}$ chitin), and luminescence was measured over time using a Tecan Infinite® M Plex (Tecan, Männedorf, Switzerland) plate reader at room temperature.

Immunoblot analysis

*N. benthamiana* leaf tissue was frozen in liquid nitrogen and ground into a fine powder. Soluble protein was extracted by adding an equal volume of lysis buffer (50 mM HEPES (pH 7.5), 1 mM EDTA, 2% PVPP, 5 mM DTT, 1 mM PMSF). The samples were mixed by rotating at 4°C for 2 minutes, prior to centrifugation at 4 °C for 15 minutes at 17 000 xg and the supernatant was collected. Approximately 20 µg of each protein solution was separated on two 15% SDS-PAGE gels. The first gel was stained with Coomassie blue to assess protein loading across samples, while proteins from the second gel were transferred onto a 0.22 µM nitrocellulose membrane. Blots were probed with HRP-conjugated mouse anti HA-tag (1:2000 dilution) either from ABclonal (Woburn, Massachusetts) or for the CtNUDIX effectors from Roche (Basel, Switzerland) due to low levels of CtNUDIX protein accumulation. Pierce™ ECL substrate (Thermo Fisher Scientific, Waltham, Massachusetts) was added to the immunoblots and chemiluminescence detected using a ChemiDoc imager (Bio-Rad, Hercules, California).

Nudix hydrolase enzyme assays

The phosphomolybdate Nudix hydrolase enzyme assays and the mRNA decapping assays were performed as described previously (*30*). To assess inositol pyrophosphohydrolase activity 5-PP-$InsP_5$ was first synthesized from $InsP_6$ (Merck, Darmstadt, Germany) in a 500 µL reaction volume following previously described methods (*63*) using the purified *Eh*IP6KA protein. The resulting 5-PP-$InsP_5$ was purified using the previously described gel electrophoresis-based method (*64*). Purified 5-PP-$InsP_5$ was incubated with 5 µM recombinant protein in 50 mM Tri-HCl pH 8.0, 5 mM $MgCl_2$ at 37 °C for 60 minutes. After incubation the reaction products were separated and identified using previously described methods (*64*) with minor modifications, we utilized a smaller gel (8.3 x 7.3 x 0.1 cm) and ran the gel at 300V for approximately 5 hours, until the dye front was 2/3 of the way through the gel.

Protein crystal structure determination

Crystallization screening with purified MoNUDIX protein (amino acids 35 to 156) was conducted using a Mosquito robot (STP LabTech, Melbourn, UK) in a 96-well plate format using sparse matrix screens. The sitting drop vapor-diffusion method of crystallization was used and drops consisting of 100 nL 30 mg/mL MoNUDIX containing 18 mM $InsP_6$ combined with 100 nL reservoir solution were equilibrated against a 100 µL reservoir solution. The reservoir solution resulting in the MoNUDIX crystals analyzed in the study was 200 mM Potassium thiocyanate with 20% PEG3350 from the SG1™ Screen (Molecular Dimensions, Newmarket, United Kingdom). To create the cryoprotectant 80 µL of the reservoir solution was combined with 10 µL of glycerol and 10 µL of ethylene glycol. The crystal was transferred to the cryoprotectant before flash cooling in liquid nitrogen. Diffraction datasets were collected on the MX2 beamline at the Australian Synchrotron (*65*). The highest resolution dataset allowed by the beamline geometry was selected for processing in XDS, and then scaled using AIMLESS in the CCP4 suite (*66*, *67*). The MoNUDIX crystal structure was determined using maximum-likelihood molecular replacement (MR) with Phaser in Phenix (*68*). The MR search model was an AlphaFold (*69*) model of the MoNUDIX sequence used for crystallization (sequence in **Table S2**). For all datasets automated model building and initial refinement was completed using Phenix AutoBuild (*70*). Subsequent model building was carried out manually in Coot (*71*) in-between rounds of automated refinement using Phenix Refine (*72*). Analysis of the final structures was performed with Coot (*71*), ChimeraX (*73*), and ConSurf (*74*) (default parameters were used for analysis). Data collection and refinement statistics are listed in **Table S5**. Map coordinates and structure files have been deposited in the Protein Data Bank under ID 8SXS.

Micro-scale thermophoresis (MST)

MST experiments were performed on a Monolith NT.115 instrument (NanoTemper Technologies, Munich, Germany) at 25 °C. MoNUDIX and $MoNUDIX^{KKEE}$ were labelled with Alexa Flour 647 succinimidyl ester (Thermo Fisher Scientific) and used at a final concentration of 50 nM and 110 nM, respectively. In brief, 20 µM of the protein solution was incubated with 2-fold molar excess of the fluorophore for 2 hours in the dark at room temperature, and the free dye was subsequently removed using a PD-10 desalting column (Cytiva). A stock solution of $InsP_6$ was serially diluted in buffer (10 mM HEPES pH 8.0, 150 mM NaCl), mixed 1:1 with the labelled protein, and loaded into standard capillaries (NanoTemper Technologies). MST measurements were recorded using 20 % LED power and 20 to 80 % MST power and analyzed using MO. Affinity Analysis software 2.2.7 (NanoTemper Technologies).

RNA extractions and RT-qPCR

For *N. benthamiana* samples, at 3 days post-infiltration approx. 100 mg of leaf tissue was collected from each infiltration site and frozen in liquid nitrogen. For RNA extraction and purification, the Monarch® Total RNA Miniprep Kit (NEB, Ipswich, Massachusetts) was used following the recommended protocol for plant tissue, with tissue lysis achieved by grinding the plant tissue into a fine powder while frozen in liquid nitrogen. cDNA synthesis from the purified RNA was achieved using the LunaScript® RT SuperMix Kit (NEB). qRT-PCR was performed using the Luna® Universal qPCR Master Mix (NEB) on a ViiA 7 PCR System (Applied Biosystems, Waltham, Massachusetts), with primers listed in **Table S6**. Expression levels were calculated relative to the geometric mean (*75*) of the reference genes, *Nb*Ube35 (*76*), and *Nb*eIF1a.

For *O. sativa* and *H. vulgare* samples, extraction of total RNA from plant tissue or fungal mycelium followed by cDNA synthesis was completed as described in (*53*). Primers used are listed in **Table S6** and marked with the suffix "qPCR". For the determination of transcript abundance or gene copy number, the protocols were similar to (*53*).

Betalain extraction and measurement

We used a modified version of a previously described betalain extraction method from *N. benthamiana* leaves (*77*). In brief, at 3 dpi six 4 mm leaf disks were excised from the infiltrated area, incubated for 30 minutes with 30 rpm rotation in 1 mL 50% methanol, and then 200 µL from each solution was transferred into a transparent 96-well plate and absorbance at 530 nm was measured. All values were zeroed using the absorbance measurement of 50% methanol alone.

Generation of RNAi strains of *M. oryzae*

The RNAi cassette from plasmid pRedi (*78*) was used to generate an RNAi construct targeting *MoNUDIX* transcripts. The 312-bp sense and antisense fragments were amplified from genomic DNA of *M. oryzae*, using the primers RNAi(Nudix1)-fw and RNAi(Nudix1)-Rv, and RNAi(Nudix1)i-fw and RNAi(Nudix1)i-Rv, respectively (sequences listed in **Table S6**). The sense and antisense fragments were used to replace the XhoI-SnaBI and BglII-ApaI fragments of pRedi and were thus separated by 135 bp of the intron of the *M. oryzae* Cut2 gene (NCBI accession: XM_365241.1), existing in pRedi, as a linker (*78*). Plasmid inserts were subsequently sequenced to verify sequence accuracy. The resulting 6.0-kb RNAi construct was excised from pRedi by DraI digestion, purified by gel elution, and transformed into conidial protoplasts of M. oryzae, and single spore isolates were generated as described previously (*79*). Knockdown of the target genes was confirmed by qRT-PCR at 28 hours post inoculation using previously described methods (*80*). At this point, rice plants showing satisfactory reduction of the transcription levels were used in our standard conidial spray inoculation and leaf sheath assays.

Analysis of *MoNUDIX* RNAi infection phenotypes

Susceptible rice variety YT-16 was used for *MoNUDIX* RNAi experiments. Rice leaf sheath inoculations were performed as described (*81*) with the following modification. We used sheath pieces that were thinner trimmed sheaths (~3 cell layers thick). Briefly, 7-cm long leaf sheath pieces from 3-week-old plants were placed in a sealable Pyrex glass moist chamber. Leaf sheath sections were placed on inverted 8-well PCR tube strips to avoid contact with wet paper and to hold the epidermal cells directly above the mid-vein horizontally flat for the uniform distribution of inoculum in trimmed leaf sheath pieces (*82*). A spore suspension ($10^4$ spores/mL in sterile 0.25% gelatin) was prepared from 10-day-old cultures and injected into one end of the sheath using a 100-µL pipette. Each segment was trimmed at 18 to 30 hpi and imaged immediately by laser confocal microscopy. Biological replicates were independent experiments performed with fungal cultures fresh out of frozen storage and with new rice plants. All conclusions are supported by at least 3 biological replicates, with each replication including observation of ~100 infection sites. Confocal imaging was performed with a Leica SP8 confocal microscope system using two water immersion objectives, C-Apochromat 40x/1.2 WCorr. and C-Apochromat 63x/1.2WCorr. Excitation/emission wavelengths were 358 nm/461 nm for cell wall fluorescence. Image acquisition and processing were done using Leica LAS X 2020 software. For ROS analysis, rice leaves inoculated with conidium suspensions ($3x10^5$ spores/mL) of the wild-type and RNAi strains were stained with DAB at 32 hpi as described previously (*81, 83*). The leaves inoculated with RNAi and wild type strains were incubated in 1 mg/mL DAB solution, pH 3.8, at room temperature for 8 hours and destained with ethanol:acetic acid solution (94:4, v/v) for 1 hour.

Deletion of *MoNUDIX*

Design of primers, gRNAs and the prediction of gene sequences was done using the *M. oryzae* 70-15 genome assembly MG8 (GCF_000002495.2) and ASM292509v1 of isolate Guy 11. All primers are listed in **Table S6**. For genome editing, protoplast transformation was performed as described in (*84, 85*). Deletion of the first *MoNUDIX* paralog was done using CRISPR/Cas9 mediated genome editing by replacing the coding sequence of *MoNUDIX* with a PCR product encoding a hygromycin resistance cassette (HygR). To enable homologous recombination, the HygR was flanked by 50 bp homologous regions as described by (*86*). The flanks were added by using primer MhNx_HygR F/R and plasmid pTK144 as a template for HygR. To exclude off-target effects, different gRNAs were used to generate three independent mutants. The mutants with a single *MoNUDIX* gene deletion were generated using gRNA-1 (ΔΔ*MoNUDIX*-M1[1] and -M2[1]), gRNA2 (ΔΔ*MoNUDIX*-M3[1], -M4[1], and -M5[1]) and gRNA-3 (ΔΔ*MoNUDIX*-M6[1] and -M7[1]), respectively. gRNA was generated with primers gRNA-1/-2/-3 similar to the procedure described in (*85*). PCR analysis was conducted to verify successful gene replacement through homologous recombination (HR). In all these mutants the second copy of the *MoNUDIX* gene was still present (fig. S2A). To generate double gene deletion mutants, three mutants with a single gene deletion were selected (ΔΔ*MoNUDIX*-M2[1], -M3[1] and -M6[1]). The second gene replacement was conducted using either fenhexamid or nourseothricin resistance cassettes (FenR/NatR) flanked by 50 bp homologous sequences, generated with primers MhNx_pTel_F/R and pTelFen/Nat as a template, see (*85*). Protoplasts of the single-deletion mutants were transformed with the FenR/NatR encoding DNA-repair-template and Cas9-RNP. The ΔΔ*MoNUDIX*-1 mutant was generated using gRNA-2 and NatR, the ΔΔ*MoNUDIX*-2 mutant using gRNA-3 and FenR, and the ΔΔ*MoNUDIX*-3 mutant using gRNA-1 and FenR. Again, PCR analysis was performed to genotype resulting mutants and the absence of both Nudix genes was verified using the gene-specific primer pair MoNX_Fl_F and MoNX_Fl_R (fig. S2B). An additional PCR analysis with one primer located within the resistance cassette and the other outside the modified region (primers: SeqMoNX_F &

pTel_F) confirmed the integration of FenR or NatR at the second gene locus. Additionally, qPCR was performed to confirm the absence of *MoNUDIX* transcripts in ΔΔ*MoNUDIX-1/-2/-3* (primers: qPCR MoNUDIX F/R and qPCR MoACTIN F/R).

Constitutive expression of *MoNUDIX*

Generation of mutants constitutively expressing *MoNUDIX:mRFP* was done as described in (*53*). The sequence of *MoNUDIX* was amplified with primers Nx_pTK Gib and Nx_pTK Gib-Stop and the gene sequence was inserted into the *Bcu*I/*Not*I linearized vector pTK144 by Gibson assembly. The expression construct was amplified using primers pTK144_OE F/R. Transformants were selected using HygR and screened by PCR for the insertion of the construct. Additionally, qPCR was performed to confirm the accumulation of transcripts of *MoNUDIX:mRFP*.

Complementation of *MoNUDIX* deletion

To verify that the observed phenotypes correlate with the deletion of *MoNUDIX* and is not caused by off targets effects potentially introduced by CRISPR/Cas9, the *MoNUDIX* locus was amplified from the genome of *Mo*WT with approximately 500 bp of downstream and 1000 bp of upstream UTR flanks (NXcompl_Gib F/R) and cloned into pTelFen linearized with *Asc*I and *BamH*I. 15 µg of the purified PCR product (generated with primers SeqMoNX_F /Nwcomp R) were used together with 1 µg pTelNat or pTelFen to enable co-selection, and a Cas9-RNP, that targets the HygR (generated with primer gRNA_Hyg), for the co-transformation of protoplasts. Insertion of the DNA repair template into the original *MoNUDIX* locus, which was replaced by HygR, was verified by PCR using primers NW-KO-NX-F/MoNX-Fl-R and NW-Comp F2/R2 (fig. S2C). Since ectopic insertion might occur, the copy number of the complemented *MoNUDIX* gene was determined by qPCR for the mutants ΔΔ*MoNUDIX*-1/2$^{comp}$, as described in (*84*), revealing two insertions for ΔΔ*MoNUDIX*-1$^{comp}$ and five insertions for mutant ΔΔ*MoNUDIX*-2$^{comp}$.

Site directed mutagenesis was performed to complement double gene-deletion mutants with potentially enzymatic inactive versions of the MoNUDIX protein. Therefore, codons encoding for amino acids within the Nudix box were replaced with alanine encoding codons (mutant 1: E83A, mutant 2: E82A&E83A, mutant 3: R78A&E79A and mutant 4: E82A&E83A&R78A&E79A). The primers used for site directed mutagenesis were: NxF-Fl_mut F/R, NX78/79_F/R,

NXKombi78/79/82/83F/R, NXFLFMUT F/R, NX82/83F/R, and NX83F/R. Constructs were amplified with overlapping sequences, ligated with Gibson-Assembly and subsequently inserted into plasmid pTelFen to fuse the construct with FenR. The resulting plasmid containing Promotor$^{NUDIX}$:*MoNUDIX*$^{mutant}$:Terminator:FenR was linearized and introduced into the gene deletion mutant ΔΔ*MoNUDIX*-1 by PEG-mediated protoplast transformation. Expression of each mutated *MoNUDIX* gene and the insertion copy number was verified by qPCR.

Co-expression of MoNUDIX:mRFP and Pwl2:GFP

For the mRFP in-locus tagging of *MoNUDIX,* 1000 bp of the promoter region, the Nudix encoding sequence with an artificial SNP in the gRNA1 target region, an mRFP encoding sequence and approx. 500 bp of the terminator sequence were amplified by PCR with overlapping primers and cloned by Gibson-Assembly into pTK144 (5UTR_NXiL_Gib F, NxiLRsnp, NxiLFsnp, NxiLRtomRFP, NxiLmRFPF, NxiLmRFPR, 3UTRF, 3UTR_Gib R). The artificial SNP was designed not to change the amino acid sequence, but prevent the repair template from Cas9-RNP cleavage. The MoNUDIX:mRFP expression construct with the original promoter and terminator (*Promoter*$^{NUDIX}$*:NUDIX*$^{SNP}$*:mRFP:Terminator*$^{NUDIX}$) was amplified and together with Cas9-gRNA1-RNP used for co-transformation of *Mo*WT. For selection, 1 µg of pTelFen was used. Fen selected transformants were screened by PCR for the insertion of the construct into at least one original *MoNUDIX* locus (Nwcomp F/ MoNX-Fl R).

Pwl2:GFP was used as a BIC marker to check for co-accumulation. To express Pwl2:GFP under the control of the original promoter, the native locus of *PWL2*, including the 1000 bp native promoter and terminator region, was amplified by PCR from *M. oryzae* isolate Guy11. Additionally, a GFP encoding sequence was amplified from plasmid pSite4NB and fused to the *PWL2* construct by Gibson Assembly (primer used: PWL Promoter F, PWL R, PWL GFP F, PWL GFP R, PWL term F, PWL term R). The construct (*Promoter*$^{PWL2}$*:PWL2:GFP:Terminator*$^{PWL2}$) was inserted into pTelFen to fuse it with the fenhexamid resistance cassette and subsequently the linearized plasmid was used for the transformation of mutants expressing MoNUDIX:mRFP under the control of the original promoter. After selection of fenhexamid-resistant transformants, the insertion of the construct was verified by PCR. Mutants showing co-expression of MoNUDIX:mRFP and Pwl2:GFP-were then inoculated on barley cultivar Ingrid (40000 conidia ml$^{-1}$).

Live-cell imaging of plants infected with *M. oryzae*

Confocal laser scanning microscopy was conducted using a Leica TCS SP8 Spectral Confocal Microscope. Inoculated barley leaves were infiltrated with Phosphate Buffered Saline (PBS) pH 7.0 (8 g $L^{-1}$ NaCl, 0.2 g $L^{-1}$ KCL, 1.4 g $L^{-1}$ $Na_2HPO_4$, 0.27g $L^{-1}$ $KH_2PO_4$) by vacuum infiltration. For mRFP detection, the samples were excited with a wavelength of 561 nm and the emission was monitored at 580–620 nm. GFP-fluorescence was captured by exciting with 488 nm and emission was monitored at 500–530 nm. To confirm mRFP fluorescence and to distinguish it from plant tissue autofluorescence, lambda scans were performed. In addition, the GFP channel was used to distinguish autofluorescence from mRFP fluorescence. Plasmolysis of the rice cells was conducted by vacuum infiltration of leaves with 0.5 M $KNO_3$. Treatments with brefeldin A (BFA) (Invitrogen, Waltham, Massachusetts) were performed on infected rice sheath tissue at 25–28 hpi ($2 \times 10^4$ spores/ml in 0.25% gelatin solution) using 50 µg $ml^{-1}$ BFA (0.1% DMSO) to inhibit specifically Golgi-dependent secretion of apoplastic effectors (*32*). Infected leaf sheaths treated with 0.1% DMSO were used as a control.

Leave samples used for WGA-AlexaFluor488 staining were cut into approximately 0.5 x 0.5 $cm^2$ tissue pieces and incubated in 1 M KOH for 1h at 37 °C. After the incubation, leaf samples were washed with PBS buffer pH 7.4 containing 0.1% TritonX. The leave samples were incubated in staining solution overnight (20 µg/ml WGA-AlexaFluor488, 50 µg/ml propidium iodide, 20 µg/ml BSA and 0.1% TritonX in PBS buffer pH 7.4). Before microscopy, the samples were washed with 1x PBS buffer pH 7.4.

Statistical analysis

Datasets with normal distributions were analysed using an independent-samples t-test (for pairs) or a one-way ANOVA with Tukey's post-hoc test for multiple comparisons. For other datasets, significance testing was completed using either a Mann-Whitney U test, when making pair-wise comparisons, or a Kruskal-Wallis H test with Dunn's post-hoc when making three or more comparisons simultaneously. The number of replicates for each experiment is stated in the appropriate figure caption. All statistical tests were completed using the 'stats' module in SciPy (*87*).

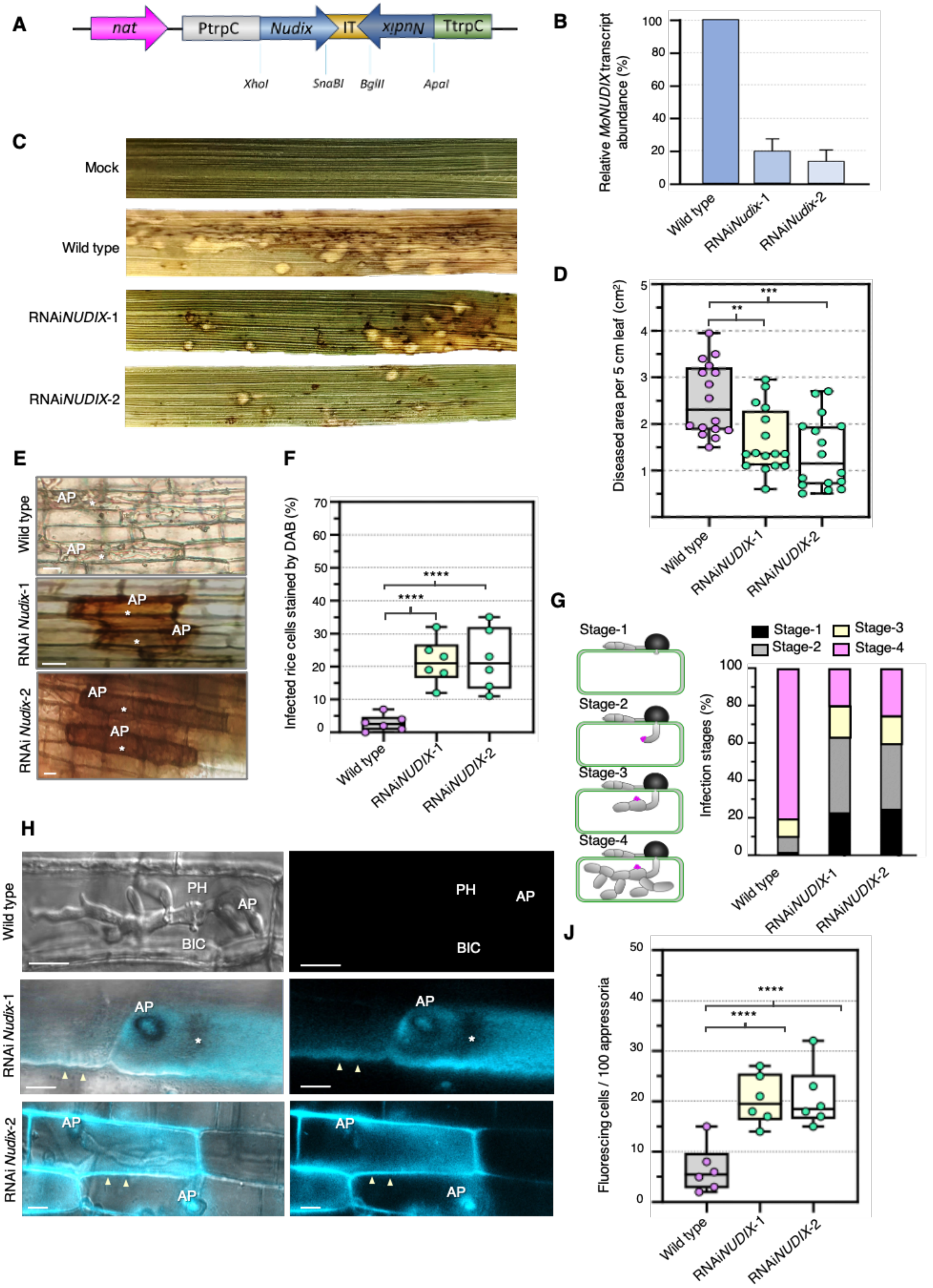
A
nat
PtrpC
Nudix
IT
Nudix
TtrpC
XhoI
SnaBI
BglII
ApaI
B
Relative MoNUDIX transcript abundance (%)
Wild type
RNAiNudix-1
RNAiNudix-2
C
Mock
Wild type
RNAiNUDIX-1
RNAiNUDIX-2
D
Diseased area per 5 cm leaf (cm²)
Wild type
RNAiNUDIX-1
RNAiNUDIX-2
E
Wild type
RNAi Nudix-1
RNAi Nudix-2
AP
F
Infected rice cells stained by DAB (%)
Wild type
RNAiNUDIX-1
RNAiNUDIX-2
G
Stage-1
Stage-2
Stage-3
Stage-4
Infection stages (%)
Wild type
RNAiNUDIX-1
RNAiNUDIX-2
H
Wild type
RNAi Nudix-1
RNAi Nudix-2
PH
AP
BIC
J
Fluorescing cells / 100 appressoria
Wild type
RNAiNUDIX-1
RNAiNUDIX-2

**Fig. S1. Silencing of *MoNUDIX* in *M. oryzae* indicates an important role in pathogen virulence and host immune suppression.** (**A**) The RNAi cassette transformed into *M. oryzae*. Consisting of the TrpC promoter (PtrpC) from *A. nidulans*, followed by 300-bp of the sense and antisense of *MoNUDIX* sequence (Nudix) separated by the Intron2 from the Cutinase2 of *M. oryzae* (IT), followed by the TrpC terminator (TtrpC). The neomycin phosphotransferase (npt) resistance cassette was used as a resistance marker. Not to scale. (**B**) Mean ± SD (n = 3) relative transcript abundance of *MoNUDIX* in infected leaf sheaths at 28 hpi (hours-post inoculation) with either wild-type *M. oryzae* or the RNAi strains. (**C**) Whole plant spray inoculation assays demonstrates that silencing *MoNUDIX* results in fewer, smaller lesions. (**D**) Quantification of the diseased area caused by wild type, RNAiNudix-1, and RNAiNudix-2 *M. oryzae* in 5 cm leaf segments of rice. Box and whisker plots with individual data points are shown; independent samples t-test **P=0.0018, ***P=0.0002; n = 16. (**E**) DAB staining of penetrated plant cells at 32 hpi. White asterisks indicate infected rice cells, AP indicates appressoria. Scale bars = 10 µm. (**F**) Percentages of infected cells stained by DAB. For each of the six replicates three sets of 100 cells were measured. ****P < 0.0001, independent samples t-test. (**G**) Quantification of four infection stages suggests a reduction of fungal virulence and colonization rate. For three replicates 100 infection sites were counted each. (**H**) Both wild-type and RNAi strains differentiated melanized appressoria (labelled AP) and invaded intact rice leaves, but the RNAi strains caused whole-cell (white asterisks) or cell wall fluorescence (white arrowheads) in rice under UV light, indicative of increased phenolic compound production and deposition in the cell wall. Scale bars = 10 µm. (**J**) Quantification of fluorescing rice cells decorated with single appressoria of the wild-type or RNAi strains. For each of the six replicates three sets of 100 cells were measured. ****P < 0.0001, independent samples t-test.

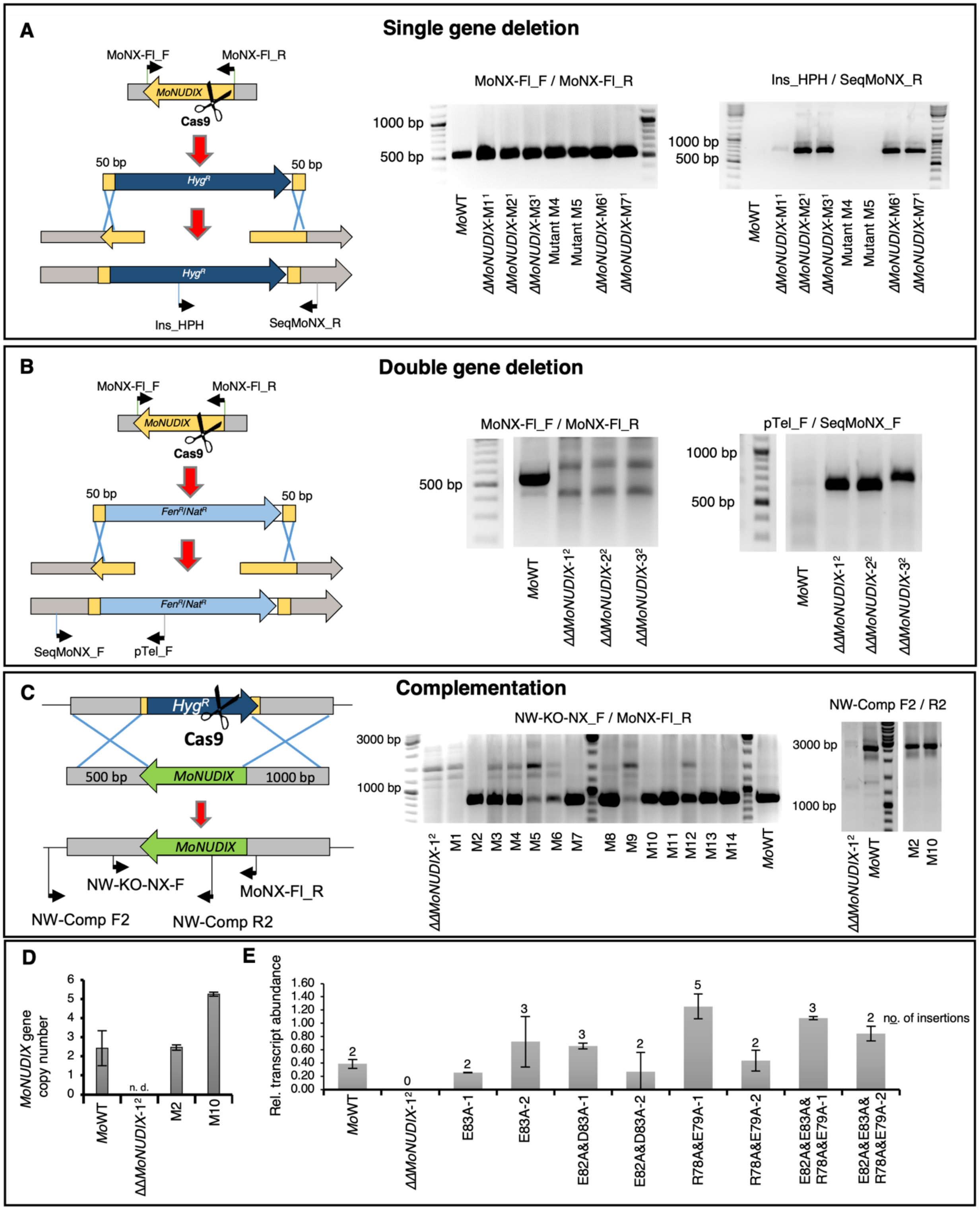


**Fig. S2. Gene deletion and complementation of *MoNUDIX* in *M. oryzae*.**

(**A**) Deletion of the first *MoNUDIX* allele was completed using CRISPR/Cas9 mediated genome editing via replacement with a hygromycin resistance encoding gene flanked by 50 bp homologous regions. Mutants M1 and M2 were generated using gRNA-1, mutants M3, M4, and M5 using gRNA-2, and mutants M6 and M7 using gRNA-3. PCR analysis using primers Ins_HPH and SeqMoNx_R was conducted to verify successful gene replacement through homologous recombination (HR). The expected size of the PCR product after HR was 686 bp. In all mutants (M1 to M7) the paralogue gene was detected by PCR, using the *MoNUDIX* specific primers MoNX-Fl_F and MoNX-Fl_R, resulting in a product of 530 bp. (**B**) Mutants M2, M3 and M6 with the single deletion were then taken to introduce the second deletion using the same approach as described in (**A**). Again, PCR analysis was performed to genotype resulting mutants and the absence of both Nudix genes was determined using the gene-specific primer pair MoNX-Fl_F and MoNX-Fl_R. An additional PCR analysis with one primer located within the resistance cassette and the other outside the modified region (primers: SeqMoNX_F and pTel_F) confirmed the integration of FenR or NatR at the second gene locus. The expected size of the PCR product after HR was approximately 670 bp while it was approximately 150 bp larger in case of NHEJ. The ΔΔ*MoNUDIX*-1 mutant was generated using gRNA-2 and NatR, the ΔΔ*MoNUDIX*-2 mutant using gRNA-3 and FenR, and the ΔΔ*MoNUDIX*-3 mutant using gRNA-1 and FenR. (**C**) For complementation, ΔΔ*MoNUDIX*-1 and -2 gene deletion mutants were co-transformed with a CRISPR/Cas9-RNP, targeting the hygromycin resistance cassette, and with a DNA repair template encoding the sequence of the *MoNUDIX* locus. The plasmid pTEL-Fen was used for co-selection of ΔΔ*MoNUDIX*-1 and pTEL-Nat for co-selection of ΔΔ*MoNUDIX*-2 mutants. To promote homologous recombination into the target locus, 500 and 1000 bp homologous flanks were used. Insertion of the DNA repair template into the original *MoNUDIX* locus was determined by PCR with primers NW-KO-NX_F and MoNX-Fl_R. If complementation was successful, a PCR product of approximately 840 bp was expected. PCR with primers NW-Comp F2 /R2 indicates an insertion of the DNA repair template into at least one *MoNUDIX* locus (PCR product size 2950 bp); mutants M1-M7 were generated from gene deletion mutant ΔΔ*MoNUDIX*-1 and mutants M8-M14 from gene deletion mutant ΔΔ*MoNUDIX*-2. (**D**) *MoNUDIX* gene copy number determined by qPCR for *Mo*WT, ΔΔ*MoNUDIX*-1 and M2/M10 (ΔΔ*MoNUDIX*-1/$2^{comp}$). (**E**) To verify the expression of *MoNUDIX* in the complemented ΔΔ*MoNUDIX M. orzyae* used for the infection experiments in Fig. 1F, transcript abundance of the mutated *MoNUDIX* genes was quantified by qPCR at 72 hpi. The *MoNUDIX* gene insertion number was calculated by qPCR as described in (*84*) and is indicated above each bar.

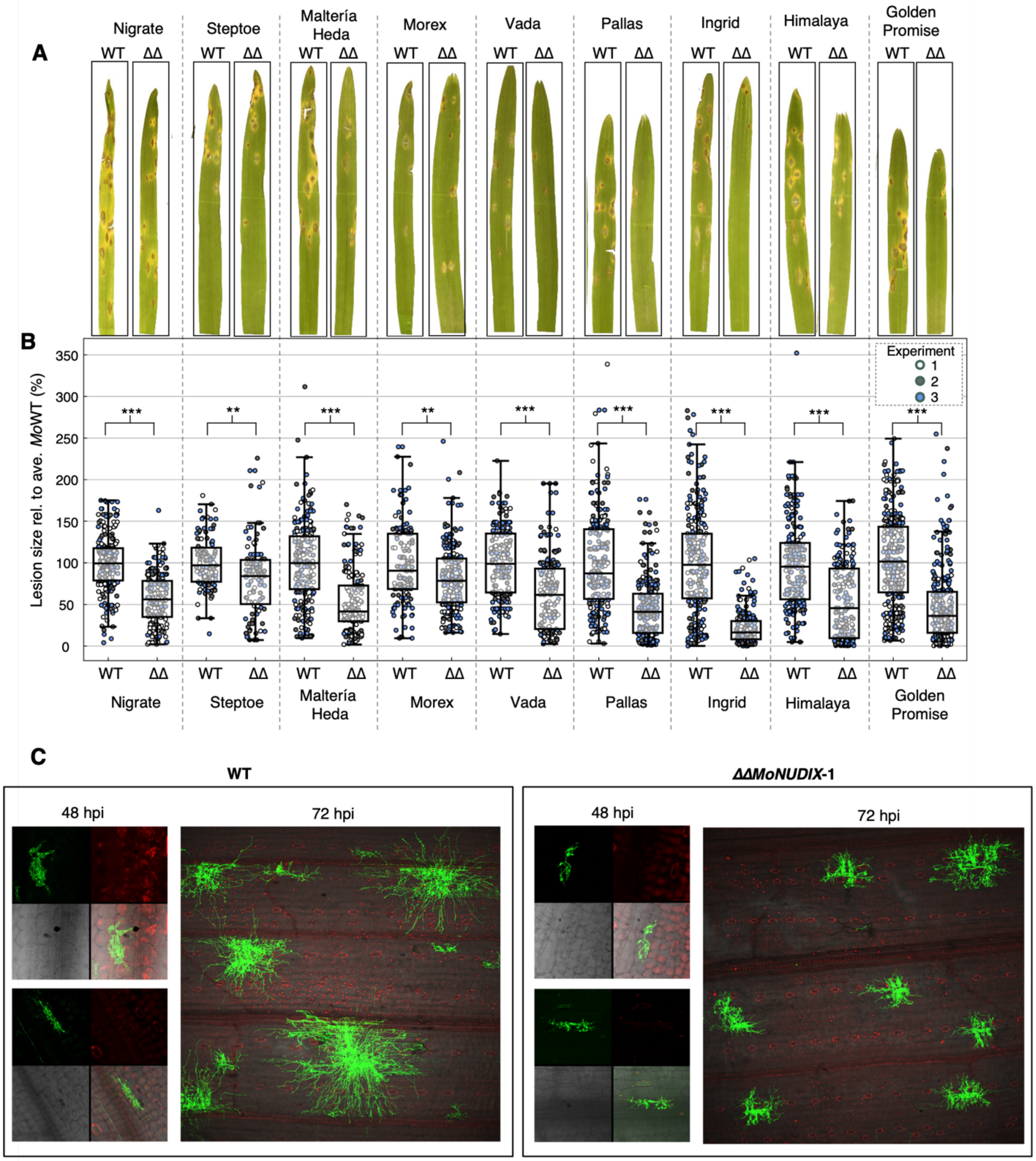


**Fig. S3.** ***MoNUDIX*** **significantly contributes to the virulence of** ***M. oryzae*** **on multiple barley cultivars.**

(**A**) Seven-day-old primary leaves of the barley cultivars Nigrate, Steptoe, Malteria Heda, Morex, Vada, Pallas, Ingrid, Himalaya, and Golden Promise were inoculated with conidia (10000 conidia $mL^{-1}$) of the *M. oryzae* wild-type isolate Guy11 (WT) and the *MoNUDIX* double gene deletion mutant ΔΔ*MoNUDIX*-1 (ΔΔ). Leaves were photographed seven days post inoculation. Representative pictures of leaves are shown for each interaction. **(B)** Blast lesion size was determined in pixels and plotted as boxplots. To combine the results of the three independent experiments, the average lesion size of WT was set to 100 %, all other values were set relative to WT. Differences between WT and ΔΔ are indicated with asterisks, as determined with the Mann-Whitney U test, $^{**}P < 0.005$, $^{***}P < 0.0005$. **(C)** Conidia of WT and ΔΔ*MoNUDIX*-1 were inoculated on barley leaves of the cultivar Ingrid and fungal hypha were stained with wheat germ agglutinin (WGA) conjugated to Alexa® Fluor 488 at 48 and 72 hours post infection (hpi). To visualize plant cells, ethidium bromide was used. Confocal laser-scanning microscopy was carried out to show the progress of the infection on barley. Each infection site was recorded using different light excitations shown as a composite figure (48 hpi) or a merged image (72 hpi). For the 48 hpi composite figure: GFP-channel, 488 nm (top left), mRFP-channel, 561 nm (top right), bright field (bottom left) and merged channel (bottom right).

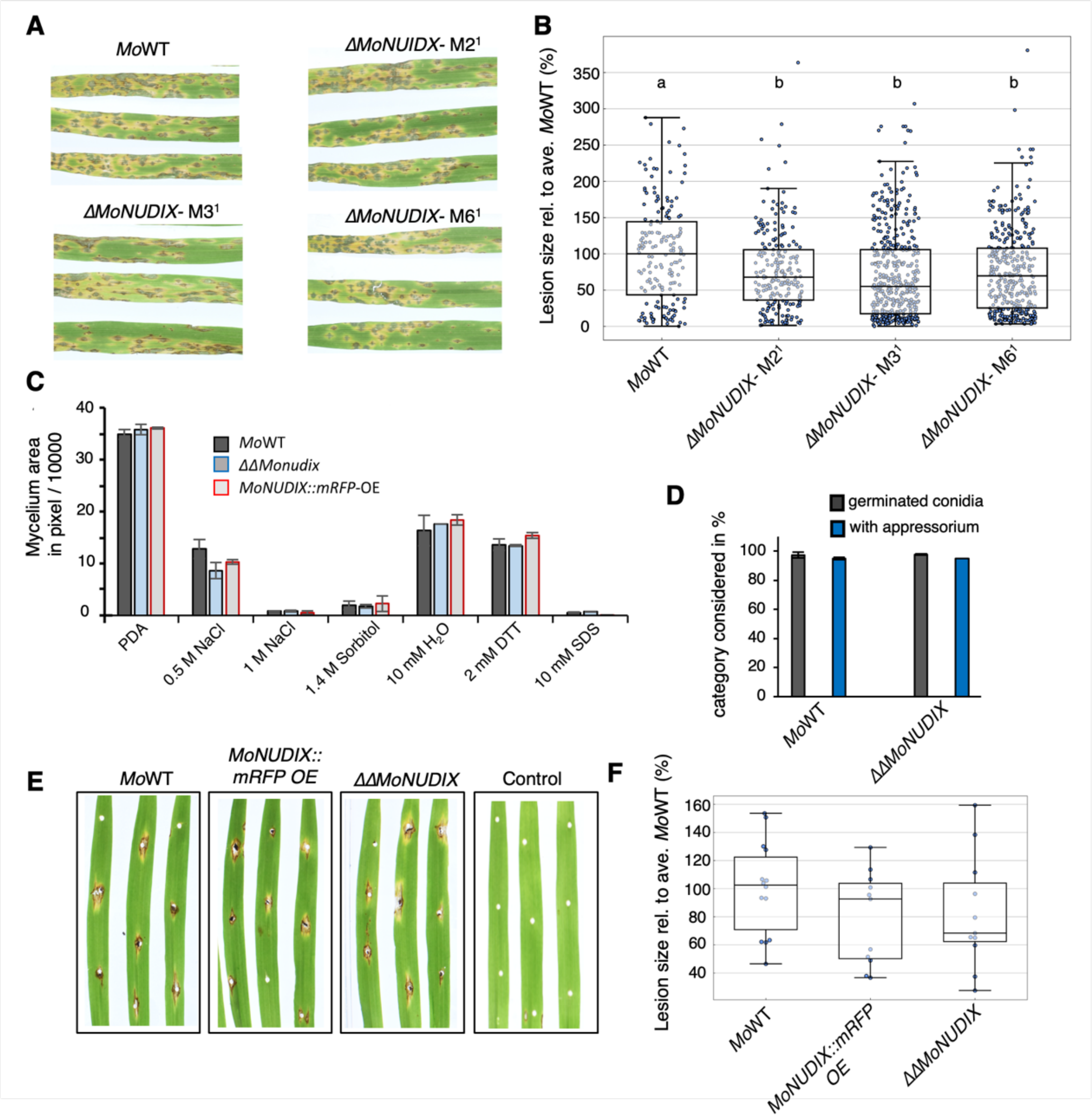


**Fig. S4. Single *MoNUDIX* gene deletion results in a very minor reduction in lesion size, and double *MoNUDIX* gene deletion specifically effects *M. oryzae* growth during infection.**
(**A**) To assess the impact of a single *MoNUDIX* gene deletion, wild-type *M. oryzae* (*Mo*WT) and mutants ΔMo*NUDIX*-M2[1], -M3[1], -M6[1] were spray inoculated on seven-day old barley leaves. Representative images of each interaction at seven days post inoculation are shown. (**B**) The size of ~ 200 lesions across 10 leaves were determined for each interaction using ImageJ and plotted in a box-plot diagram. Letters depict significant differences between treatments as determined with the Kruskal-Wallis H test followed by Dunn's post-hoc test ($P < 0.05$). (**C**) Comparison of vegetative mycelial growth of *Mo*WT, a constitutively *MoNUDIX:mRFP* expressing mutant (*MoNUDIX:mRFP-OE*), and ΔΔ*MoNUDIX*-1. An agar block overgrown with mycelium was placed onto minimal agar with various additives to induce pH, osmotic or oxidative stress as described in (*84*). The mycelium grown at 25 °C in the dark was photographed after five days and evaluated using ImageJ. The mean value of the overgrown mycelium area in pixels and the

standard deviation of three technical replicates are plotted. The experiment was repeated with comparable results. (**D**) *In vitro* germination test and appressoria formation rate of conidia suspensions from *Mo*WT and ΔΔ*MoNUDIX*-1 after 48 hours placed in a hemocytometer. (**E**) Seven-day-old primary leaves of the barley cultivar Ingrid were wounded with forceps. Vegetative mycelium of *Mo*WT, *MoNUDIX:mRFP-OE*, and ΔΔ*MoNUDIX*-1 were placed on the injured leaf site. After seven days lesions were photographed, and representative pictures of infection sites are shown. Injured but not mycelium-inoculated leaves were used as a control. (**F**) The lesion size of each infection site was determined in pixels using image analysis software ImageJ and plotted in a box-plot diagram. No significant differences were detected between *Mo*WT and ΔΔ*MoNUDIX*-1 or *MoNUDIX:mRFP-OE* (Mann-Whitney U test, $P > 0.05$).

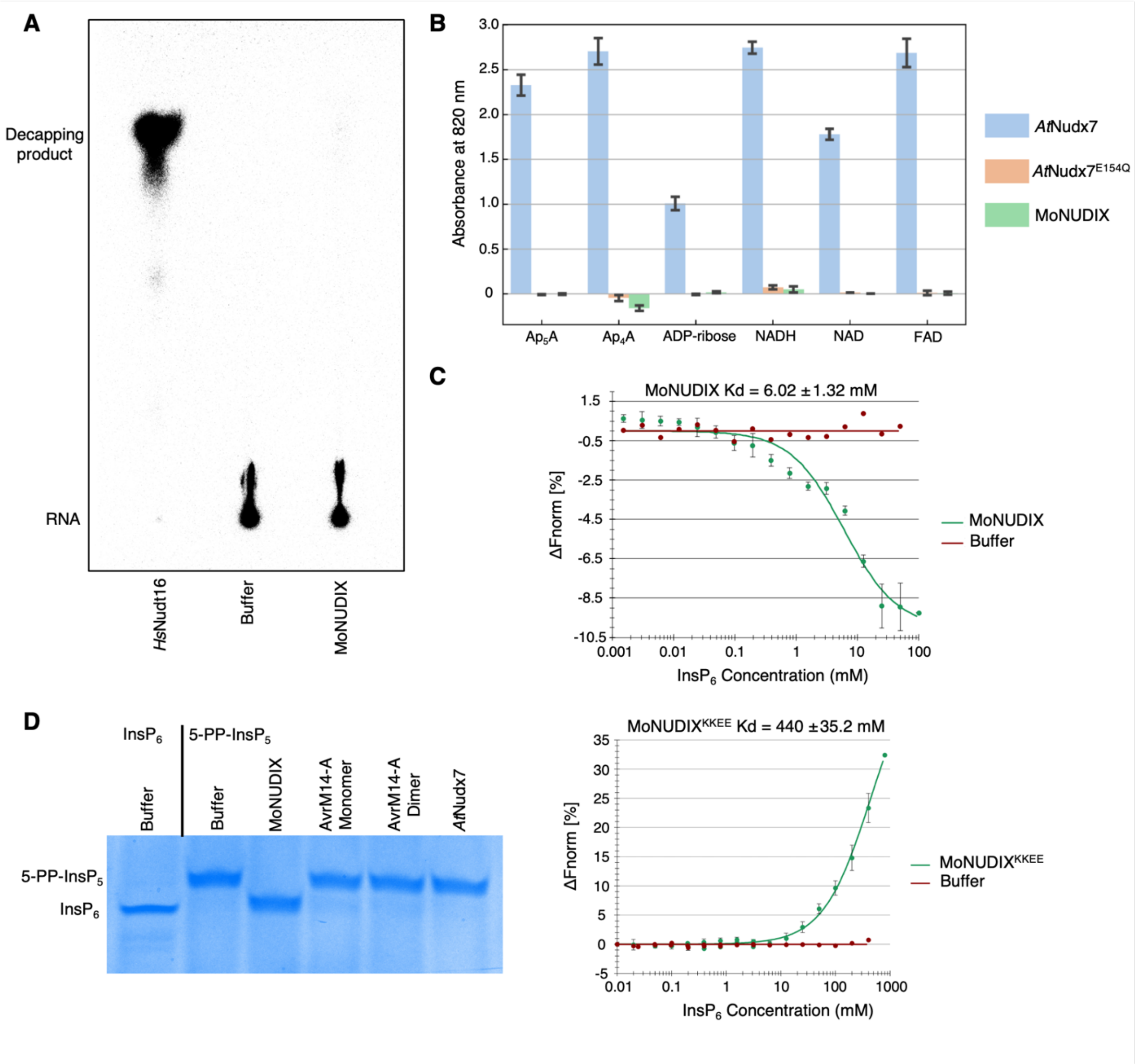


**Fig. S5. MoNUDIX specifically hydrolyses inositol pyrophosphates and active site lysines are involved in inositol polyphosphate binding.**
(**A**) Purified proteins (*Hs*Nudt16, a known mRNA decapping enzyme (*88*) and MoNUDIX) were incubated with $^{m7}Gp_{32}$pp-RNA and the reaction products analysed by thin-layer chromatography (TLC). Capped RNA remains at the origin of the TLC plate, whereas decapping products migrate up the plate. (**B**) Purified proteins (*At*Nudx7, *At*Nudx7$^{E154Q}$, and MoNUDIX) were incubated with 2 mM of the indicated substrate at 37 °C for 30 minutes. Substrate hydrolysis was detected via the production of a blue-coloured phosphomolybdate complex that absorbs light with a wavelength of 820 nm. Results are mean absorbance ± SD (n = 3). A buffer only control without any Nudix hydrolase protein was used to blank the spectrophotometer before measurement. *At*Nudx7 and *At*Nudx7$^{E154Q}$ were used as positive and negative controls, respectively. (**C**) Normalized microscale thermophoresis (MST) binding curves of MoNUDIX (top) and MoNUDIX$^{KKEE}$ (bottom) in the presence of $InsP_6$, alongside a protein storage buffer control. The binding curve yields a Kd of 6.02 ± 1.32 mM for MoNUDIX and a Kd of 440 ± 35.2 mM for MoNUDIX$^{KKEE}$.

Protein concentrations were kept constant while the $InsP_6$ concentration varied. (**D**) Purified protein (MoNUDIX, AvrM14-A monomer, AvrM14-A homodimer, *At*Nudx7) at a concentration of 5 µM was incubated with 5-PP-$InsP_5$ for 60 minutes at 37 ºC. As a control, buffer-only reactions were completed with $InsP_6$ and 5-PP-$InsP_5$. All reaction products were separated using a polyacrylamide gel and visualised by staining with toluidine blue.

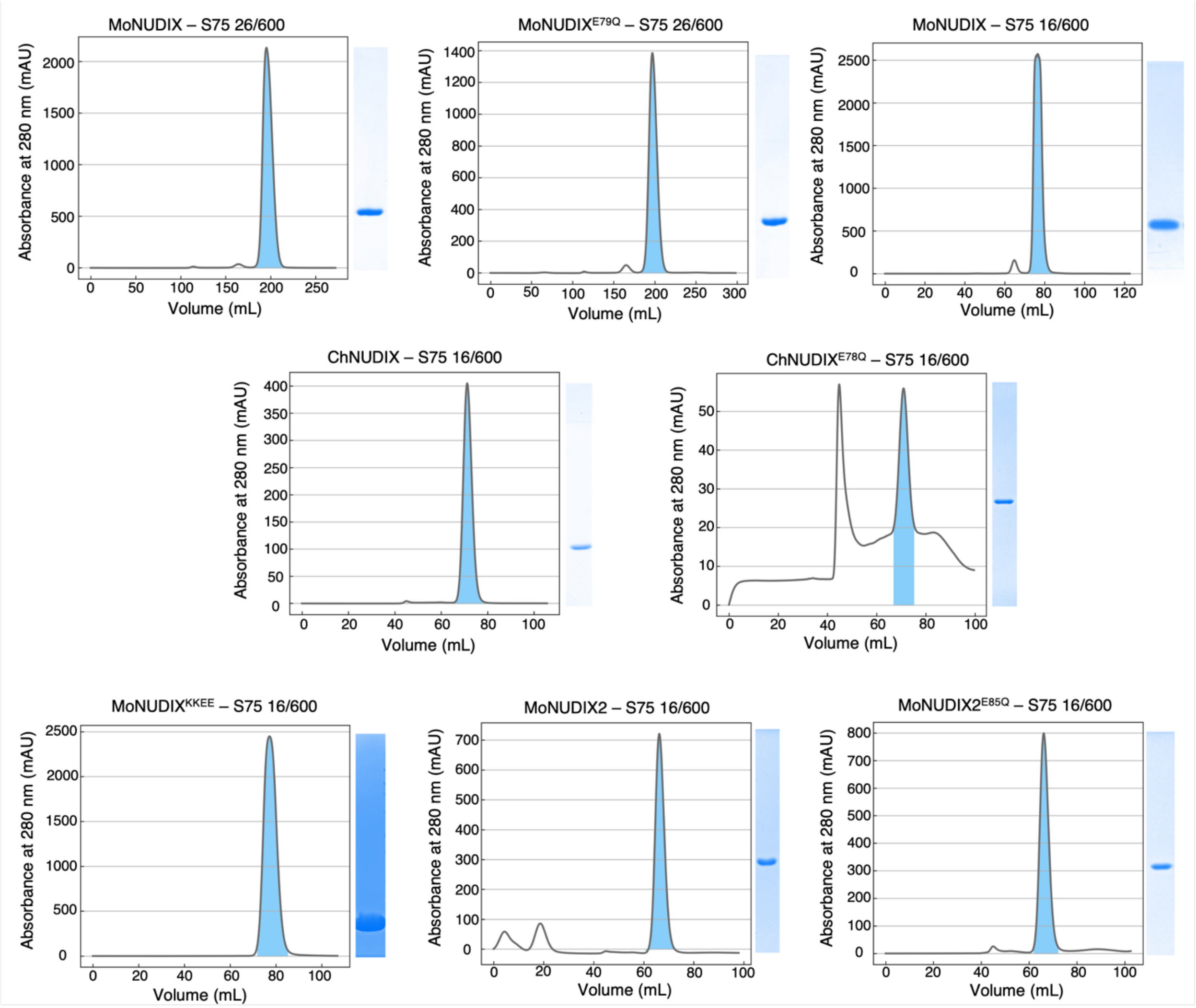


**Fig. S6. All effector proteins used in enzymatic assays were purified to homogeneity.**
Size exclusion chromatography (SEC) profiles for all effector proteins purified in this study. The label at the top of each profile indicates the protein and the Highload Superdex column (Cytiva) used in the purification process. The area under the peak shaded blue indicates the volume collected for each effector. Alongside each profile is a Coomassie-stained SDS-PAGE gel demonstrating the purity of the final protein sample collected following SEC and used in enzyme assays.

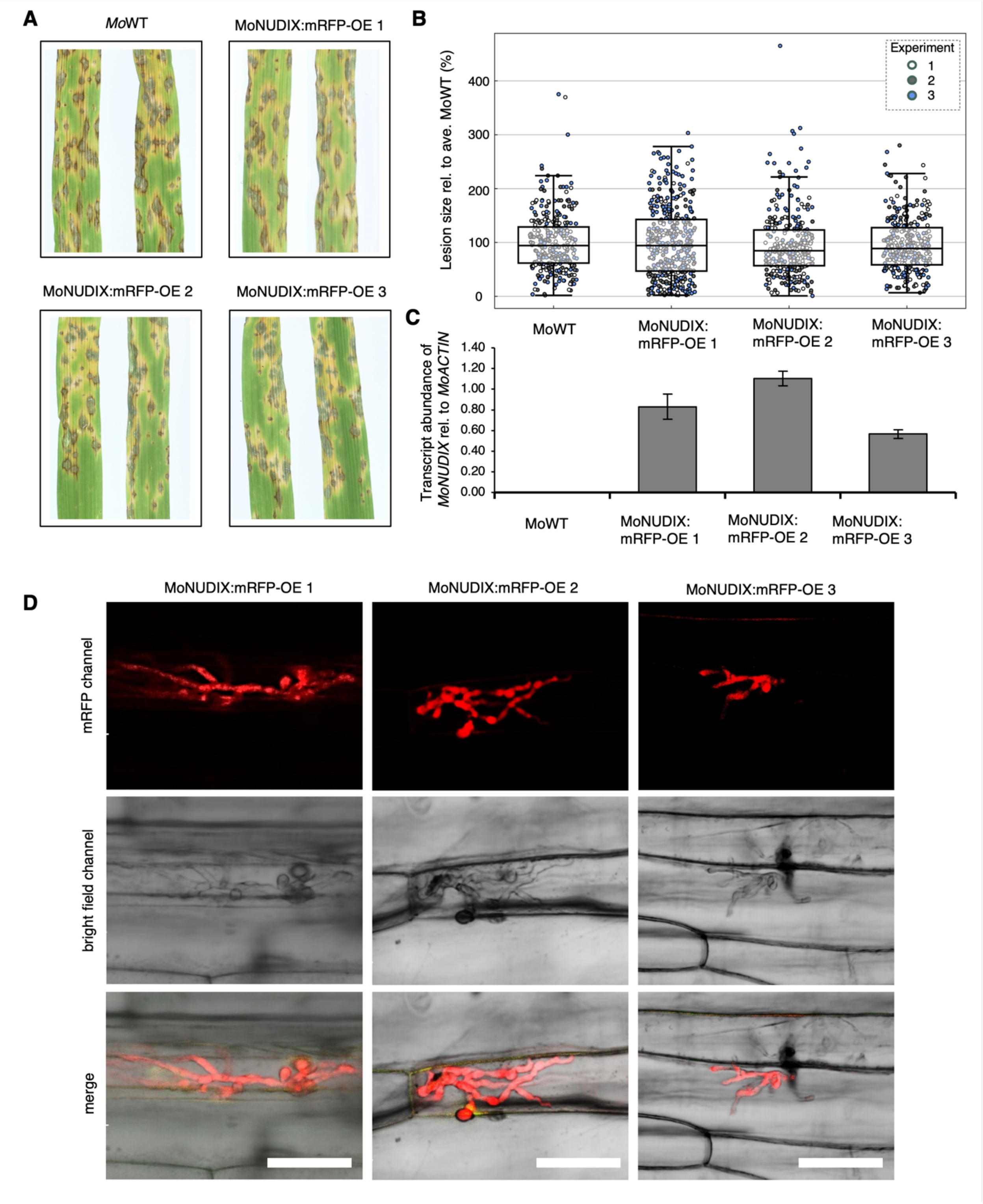


**Fig. S7: Constitutive expression of MoNUDIX:mRFP in *M. oryzae* Guy11.**
(**A**) To assess the impact of constitutive expression of MoNUDIX:mRFP on virulence, conidia of wild-type *M. oryzae* (*Mo*WT) and three mutants (MoNUDIX:mRFP-1/-2/-3) were spray inoculated on seven-day old barley leaves. Representative images of each interaction at seven days post inoculation are shown. (**B**) For each interaction, the size of ~ 100 lesions across 5 leaves for

three independent experiments was determined using ImageJ and visualized in a box-plot diagram. No significant differences between any treatments were identified (Kruskal-Wallis H test, P = 0.34). (**C**) To verify expression of MoNUDIX:mRFP, transcript abundance in hyphae grown on PDA was quantified by qPCR. (**D**) For localization, confocal laser-scanning microscopy was performed 72 hpi using all mutants, MoNUDIX:mRFP-1/-2/-3. Scale bar: 10 µm.

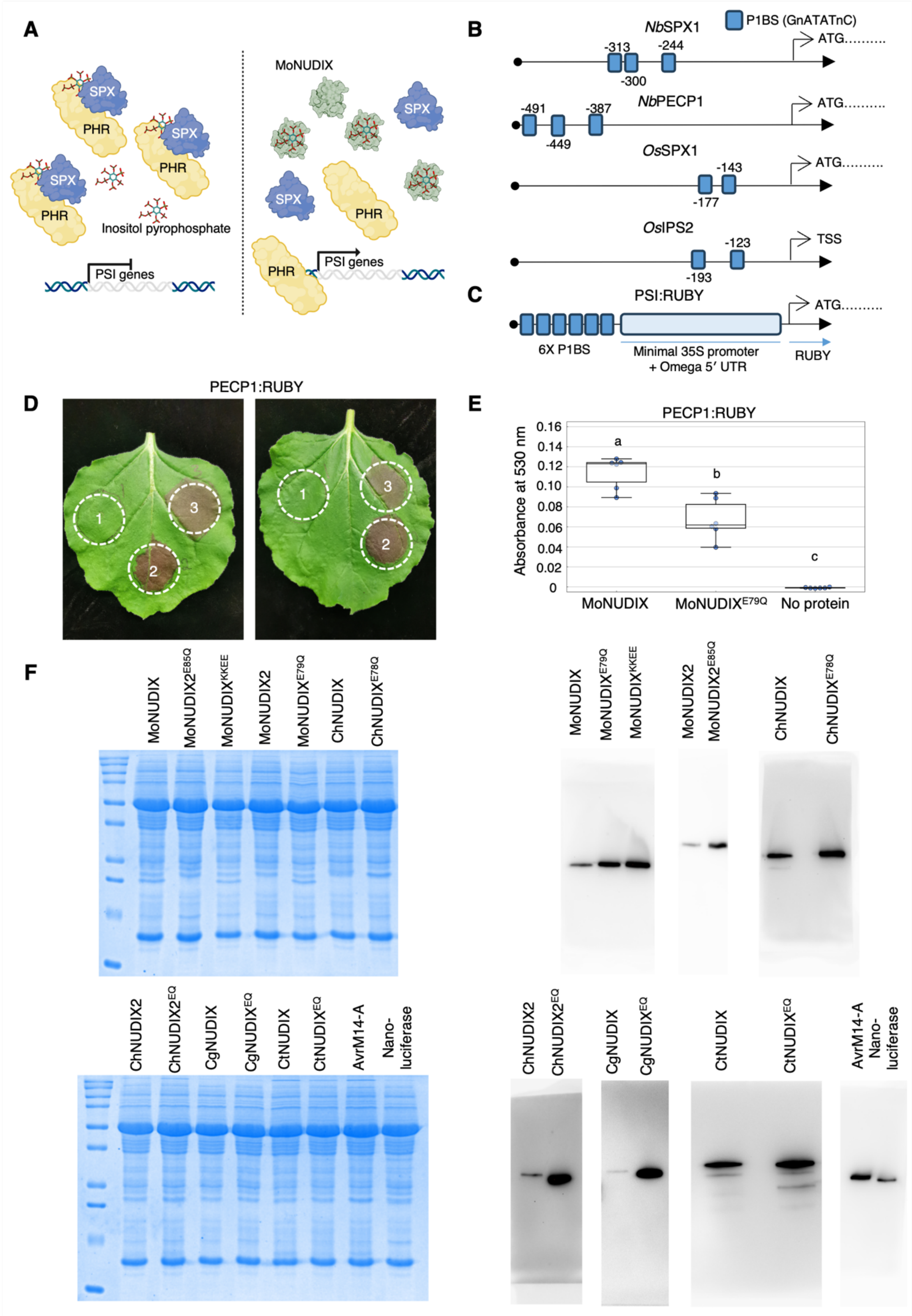


**Fig. S8. Assessing phosphate starvation induction by the Nudix effectors in *N. benthamiana***

(**A**) The hydrolysis of PP-InsP by MoNUDIX in plant cells should activate the expression of phosphate starvation induced (PSI) genes in plants by releasing PHRs from SPX proteins. (**B**) A schematic of the 500 base pairs upstream of the *Nb*SPX1, *Nb*PECP1, and *Os*SPX1 start codons with the PHR1-binding site (P1BS) elements indicated. The 500 bp promoter of *Os*IPS2 is also shown; however, as IPS2 is a long non-coding RNA and does not have a clearly defined start codon, the predicted transcriptional start site (TSS) is indicated instead. (**C**) A schematic of the PSI:RUBY promoter and 5′ UTR. The synthetic gene contains six P1BS elements in the promoter, a minimal 35 S sequence, and the Omega 5′ UTR sequence from the Tobacco Mosaic Virus. Only the start of the RUBY CDS is depicted. Not to scale. (**D**) Representative leaf images demonstrating production of the red betalain pigment in *N. benthamiana* leaf tissue co-transformed with the PEPC:RUBY promoter/reporter and wild-type MoNUDIX (labelled as 2), MoNUDIX$^{E79Q}$ (labelled as 3), or no effector (labelled as 1). (**E**) The absorbance at 530 nm of extracts from transformed *N. benthamiana* leaves. There were 6 biological replicates for each treatment, as indicated by the dots beneath the boxplots. Letters denote significant differences between treatments, as determined by a one-way ANOVA followed by Tukey's post-hoc test; $P < 0.001$. (**F**) (Left) Coomassie-stained SDS-PAGE protein gels demonstrate equivalent total protein amounts in the soluble *N. benthamiana* protein extracts from the agroinfiltrated plant tissue used for western blotting. The effector that should be present in each sample is indicated along the top of the gel; the first lane of both gels contains the Precision Plus Protein Dual Color Standards (Biorad, Hercules, California). (Right) Total protein extracts from *N. benthamiana* leaf tissue agroinfiltrated with a construct to express a HA-tagged protein, as labelled along the top of each blot, were analysed by western blotting. Blots were probed with mouse anti-HA HRP-conjugated antibodies.

**Table S1. The *Magnaporthe* and *Colletotrichum* Nudix effector family.** The names, species, host plant, protein sequences, and accession IDs of identified *Magnaporthe*, *Colletotrichum*, and *Ceratocystis* Nudix effectors.

<Table S1 in excel sheet >

**Table S2. Sequences of the purified proteins used in this study.**

<Table S2 in excel sheet >

**Table S3. Sequences of all proteins expressed in *N. benthamiana* in this study.**

<Table S3 in excel sheet >

**Table S4. Promoter sequences discussed in this study.**

<Table S4 in excel sheet >

**Table S5. Crystallography data collection and structure refinement statistics for 8SXS**

<Table S5 in excel sheet >

**Table S6. Primer sequences used throughout this study.**

<Table S6 in excel sheet >
